# ProminTools: Shedding light on proteins of unknown function in biomineralization with user friendly tools illustrated using mollusc shell matrix protein sequences

**DOI:** 10.1101/2020.03.05.978718

**Authors:** Alastair W Skeffington, Andreas Donath

## Abstract

Biominerals are crucial to the fitness of many organism and studies of the mechanisms of biomineralization are driving research into novel materials. Biomineralization is generally controlled by a matrix of organic molecules including proteins, so proteomic studies of biominerals are important for understanding biomineralization mechanisms. Many such studies identify large numbers of proteins of unknown function, which are often of low sequence complexity and biased in their amino acid composition. A lack of user-friendly tools to find patterns in such sequences and robustly analyse their statistical properties relative to the background proteome means that they are often neglected in follow-up studies. Here we present ProminTools, a user-friendly package for comparison of two sets of protein sequences in terms of their global properties and motif content. Outputs include data tables, graphical summaries in an html file and an R-script as a starting point for data-set specific visualizations. We demonstrate the utility of ProminTools using a previously published shell matrix proteome of the giant limpet *Lottia gigantea*.

## 1. Introduction

Mineralized structures are formed by many organisms across the tree of life including bacteria, molluscs, metazoans, plants and algae [1]. These biominerals are critical for fitness, playing roles in support, defence, buoyancy, regulation of ion budgets and orientation among others. Proteins have been found to be associated with many biominerals, and are hypothesised to have a key role in mineral synthesis [2–4]. In some cases the roles of such proteins is well understood. Well studies examples include the Pif protein in molluscs [5], Amelogenin from tooth enamel, Silicatein from sponge spicules and Mms6 from magnetosome synthesising bacteria [2]. However in the majority of cases the function of biomineral associated proteins remains elusive.

A common workflow in biomineralization research is to first clean a mineral preparation using detergents or oxidizing agents to remove loosely associated organic matter, and subsequently to dissolve the mineral, releasing tightly mineral-associate proteins into solution that can then be analysed using proteomic methods [6]. It is generally hypothesised that these proteins are likely to be involved in mineralization and that proximity to the site of mineralization results in their incorporation into the mineral as it grows. Some of the proteins identified may have homology to proteins of known function or recognisable domains strongly suggestive of a certain function. For example, carbonic anhydrases have been found associated with calcium carbonate minerals in several organisms [7] and may aid generation of bicarbonate as a substrate for calcification. However there are generally many proteins in such data sets which lack similarity to proteins of known function (eg [8–10]). Intriguingly, these proteins of unknown function often display unusual primary sequence characteristics, such as low complexity, biased composition and a high degree of predicted intrinsic disorder.

Informatic tools which allow biologists to easily investigate the global features of groups of proteins of unknown function relative to the background proteome are currently lacking. Thus many studies restrict their analysis of these proteins to noting the compositional biases or motifs which are obvious from manual inspection of the protein sequences. Although the human eye is good at detecting patterns, this method has the risk that important patterns in the data are missed and that rules are not applied consistently in identifying these patterns. Ideally, the context of the proteome as a whole should also be taken into account. For example, a given motif that appears several times in a protein of interest (POI) should be considered more interesting for biomineralization if it is rare in the rest of the proteome than if it is a commonly found motif.

Although there are many tools available that allow researchers to investigate the properties of protein sequences *in silico*, including analysis of compositional bias, sequence complexity, intrinsic disorder and sequence motifs, such tools are not always easy to use. Some require command line use, data input formats differ, some can only run on one protein at a time and most require post-processing of the output to format the data for statistical and graphical analyses in commonly used environments such as Microsoft^®^ Excel^®^ or R.

Here we present ProminTools, a set of easy-to-use tools for the statistical comparison of two sets of protein sequences, available as apps in the Cyverse Discovery Environment (https://de.cyverse.org/) [11]. or to run locally from a Docker^™^ container. The inputs are simply two fasta files containing the proteins of interest (POIs) and the background proteome, while the outputs include data tables and an html document containing graphical summaries of the data and interactive tables for data exploration. To demonstrate the utility of these tools, we reanalyse a published data set of shell matrix proteins (SMPs) from the giant limit *Lottia gigantea* [12].

## 2. Materials and Methods

### 2.1 ProminTools structure

ProminTools has two component programmes: “Protein Motif Finder” and “Sequence Properties Analyzer”. Both are written in Perl and R and bundled with all dependencies in Docker^TM^ (www.docker.com) containers. They can be run from Apps within the CyVerse Dicovery Environment [11] or on a personal computer via Docker^™^ Desktop. For both tools, the inputs are two fasta formatted files: one containing the protein set of interest, and the other the reference or background proteins, typically the predicted proteome of the organism of interest. The primary outputs of the tools are data tables summarising key information from the comparison of the two sequences sets. The tools use these tables to generate an html file which generates a graphical summary of the information along with explanations, statistical analyses and interactive versions of certain data tables. A publication ready SVG (Scalable Vector Graphic) formatted figure is also generated by Protein Motif Finder. The R script that generates the html file from the data tables is also an output of the tools, allowing the user to reproduce the figures in the html report and to provide a starting point for further analyses specific to the data set. For licence information for all components of ProminTools the reader is referred to Supplementary file 5.

### 2.2 Analyses performed by “Protein Motif Finder”

#### 2.2.1 Motif finding with motif-x

Protein Motif Finder uses the motif-x engine [13, 14] for motif finding. This engine was chosen because it breaks sequences down into their constituent motifs, by an iterative procedure that avoids oversimplification of motifs and prioritises motifs that are most enriched relative to a background sequence set. In this work it was always run with the recommended, conservative, binomial *p*-value of 10^−6^, but this parameter is user customisable in Protein Motif Finder. The motif width is also user customisable, while the minimum occurrences parameter is hard coded at a value of five. Motif-x is run via the R module rmotif-x, centred on each amino acid sequentially and the results are combined. This procedure means that some motifs are likely to be redundant. For example, if the central residue is ‘S’ and the motif width 7 then the motifs ‘…S.S..’ and ‘..S.S… (where dot represents any amino acid) may both be identified, but these would be collapsed to the single motif ‘S.S’. Note that this procedure is conservative with respect to the original p-value calculated by motif-x. Significant motifs are then enumerated in the POI and the background sequence sets, and motif counts and enrichments reported in the output tables. Downstream analyses do not rely on the motif-x p-value, but only on calculated enrichment values for the motifs.

#### 2.2.2 Graphical representations of motif data

To provide an visual summary of the motif data, the motifs are represented in three wordclouds in the Protein Motif Finder output, which take into account two distinct measures of ‘importance’. The first is the number of proteins in which that motif is found. The more proteins containing the motif, the more likely it is to have general importance in the function of the group of proteins. The second measure is the enrichment of the motif. The more enriched the motif the more unusual it is and thus is more likely to be involved in the specific function of these sets of proteins. A third measure attempts to combine the previous two by scaling them equivalently and then taking the product of the scaled values (PS-value). This measure prioritises motifs that are both highly enriched and found in a high proportion of the proteins.

In the output, the proteins are also clustered based on their motif number and motif enrichment. The distance measure for clustering was calculated as one minus the Pearson’s correlation coefficient for all pairwise combinations of proteins or motifs, since this method is especially robust to outliers. Hierarchical clustering was performed using the Ward.D method.

### 2.3 Analyses performed by “Sequence Properties Analyzer”

Sequence Properties Analyzer performs the following analyses:

#### 2.3.1 Amino acid enrichment

Compositional bias is analysed using fLPS [15] and the results collated to several files described in the html output of the program.

#### 2.3.2 Significance of sequence bias

To estimate the probability of obtaining the observed bias in amino acid composition in the POI set by random sampling of the background proteome, the following procedure was implemented. First the degree of bias was quantified by calculating a bias index (BI):

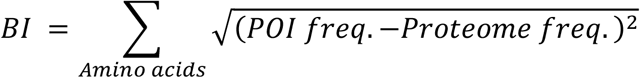

The *BI* is calculated for 1000 random samples of the background sequence set, each containing the same number of sequences as the POI set. A kernel density estimate of the distribution of *BI* is calculated and a function approximating this distribution generated. The area under the curve greater than the *BI* value of the POI set is used as an estimate of the probability of obtaining a sequence set of this degree of bias by chance, given this particular background proteome.

#### 2.3.3 Sequence complexity

The program SEG [16] is used to identify low complexity regions in the datasets using default parameters, although these are customisable by the user in Sequence Properties Analyzer. For each protein, the percentage of the seqquence identified as low complexity is calculated and a Wilcoxon rank sum test with continuity correction is used to test whether there is a significant difference in the distribution of this percentage length between the POI and the background sequence set.

#### 2.3.4 Intrinsic disorder

Predicted intrinsic disorder was calculated using the VSL2 predictor [17], due to its speed and good accuracy [18]. This is the most time consuming step of Protein Sequence Analyzer and is thus parallelized in the implementation. For each protein, the percentage of the sequence identified as intrinsically disordered is calculated and a Wilcoxon rank sum test with continuity correction is used to test whether there is a significant difference in the distribution of this percentage length between the POI and the background sequence set.

#### 2.3.5 Charged clusters

Clusters of charged amino acids are identified using the SAPS software [19].

### 2.4 *L. gigantea* shell matrix proteome data

To illustrate the utility of the ProminTools package, we used the shell matrix proteome of the giant limpet, *L. gigantea* as published by Mann *et al.* (2014) [12] which is a reanalysis of their original data [20]. The protein identifiers were extracted from table S1 of [12] and the protein sequences extracted from files Lotgi1_GeneModels_AllModels_20070424_aa.fasta and Lotgi1_GeneModels_FilteredModels1_aa.fasta which were downloaded from the JGI (https://mycocosm.jgi.doe.gov/Lotgi1/Lotgi1.home.html) on the 5/02/2020. The final set of proteins consisted of 381 sequences. This is one less than the number of accepted identifications in [12] since the protein Lotgi|172500 was not available in any database.

### 2.5 Analyses of *L. gigantea* data using ProminTools

ProminTools was run locally using the shell matrix proteins as the foreground sequences and the ‘Lotgi1_GeneModels_FilteredModels1_aa.fasta’ file as the background proteome. The filtered models were chosen as they were considered likely to be a closer representation of the true proteome of *L. giga*ntea than the ‘All Models’ set, and thus the more appropriate set for statistical comparisons.

### 2.4 Additional analyses

Proteins were clustered based on motif content as an output of the Protein Motif Finder tool. To determine an optimal cluster number, manual inspection of a plot of the cindex [21] for cluster sizes 2 – 50 was carried out. A cluster number of 24 seemed appropriate for the present work, since it captured the major patterns in the data without becoming too granular. These clusters were the input for further runs of Protein Motif Finder.

Sequence similarity was quantified using and all vs all pairwise blastp analysis, reporting the percentage identity of the top scoring high scoring pair (HSP), after applying an e-value cut-off of 0.01 and a cut-off specifying that the HSP alignment length must be at least 20% of the query length.

## 3. Results

### 3.1 ProminTools provides a user-friendly method to analyse biomineralization proteomes

The Docker^TM^ image containing ProminTools can be run via a GUI on the Cyverse Discovery Environment without any need for use of the command line. Runtimes on Cyverse are variable due to variable resource availability, but a typical analysis with either Protein Motif Finder or Sequence Properties Analyzer takes between 30 and 120 minutes to complete. Although ProminTools is designed to run in a Unix environment, it can also be run on a windows PC via Docker^TM^ Desktop with simple commands in Windows^®^ Power Shell^™^ (for details see https://github.com/skeffington/Promin-tools). On a Window^®^ 10 machine with an Intel^®^ Core^™^ i7-2600 3.4 GHz processer and 16 GB RAM, Protien Motif Finder completed analysing the *L. gigantea* data set in 11 min 8 s provided with 1 core and 2.5 GB RAM, while the Sequence Properties Analyser completed in 30 min 8 s provided with 5 cores and 5 GB RAM. The ProminTools workflow is summarised in Figure 1.

**Figure 1.**
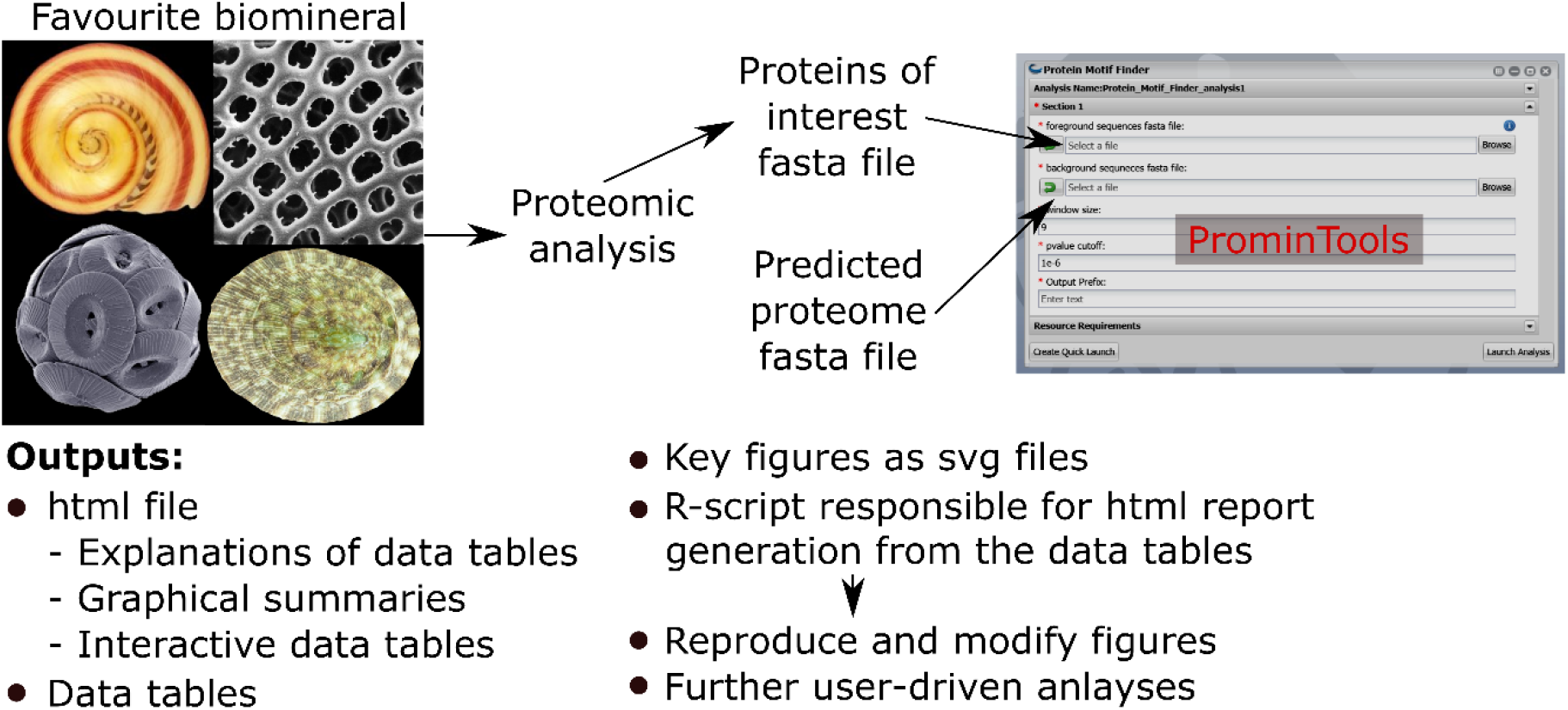
Summary of ProminTools. Proteomic datasets derived from analysis of biomineralizing organisms result in a fasta file containg the POI set which, along with the fasta file of the background proteome, make up the inputs for the ProminTools. The graphical interface shown is from the Cyverse Discovery Envronment (reproduced with permission). The outputs of the tools are detailed at the bottom of the figure. Attributions for the biomineral images are as follows: top left - *Vittina waigiensis* by H. Zell (licence: CC BY-SA 3.0), top right - Radiolarian skeleton by Hannes Grobe (licence: CC BY 3.0), bottom left - *Coccolithus pelagicus* by Richard Lampitt and Jeremy Young, The Natural History Museum, London (licence: CC BY 2.5), bottom right - *Lottia mesoleuca* by H. Zell (licence: CC BY-SA 3.0).

### 3.2 Global properties of *L. gigantea* shell matrix proteome

Previous analyses of the *L. gigantea* shell matrix proteome [12, 20, 22] had noted a tendency for the proteins to be low complexity and disordered and that some proteins were enriched in particular residues. Here, ProminTools was used to put these observations on a more quantitative footing and to discover enriched sequenced motifs in the data set from Mann *et al.* (2014) [12]. G and P rich motifs were found to be enriched most frequently among the proteins (Figure 2A). Given that we’re seeking to find the motifs that are shared within a group of proteins, Protein Motif Finder excludes motifs found in < 5% of the proteins from certain plots to prevent the picture being dominated by a highly enriched motif found in very few proteins. The result can be seen in figure 2B, where Q containing motifs displayed the greatest enrichments relative to the background proteome. For example QQP was enriched 7.5 fold while Q.N.Q was enriched 6.1 fold (see data tables in Supplementary file 2 for these numbers). In general there is often a negative correlation between the number of proteins in a set containing motifs and the enrichment of those motifs. These two measures are combined (see Materials and Methods) in Figure 2C, which emphasises motifs found in a high number of proteins with high enrichment (a high PS-value). For example, GG is found in 270 proteins and is 1.8 fold enriched, G..D in 234 proteins at 1.4 fold enrichment and NG in 249 proteins at 1.53 fold enrichment (Supplementary file 2).

**Figure 2.**
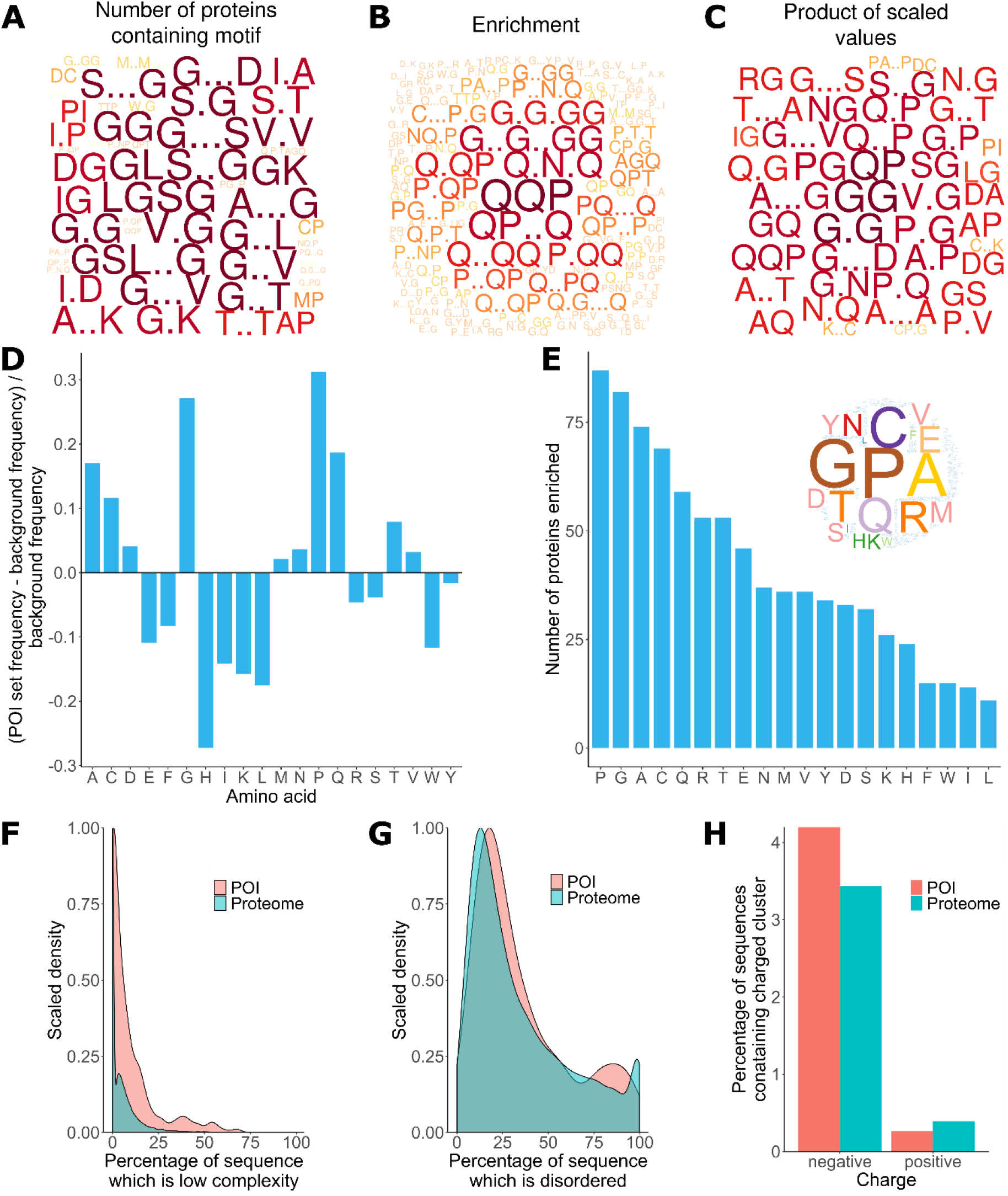
Properties of *L. gigantea* shell associated soluble proteome revealed by ProminTools. Wordclouds are displayed where the height of the letter is proportional to: **A** the number of proteins in the SMP set containing the motifs, **B** the enrichment of the motif relative to the background proteome, and **C** the product of protein number and enrichment after scaling (PS-value). **D** The enrichment of amino acids in the SMP set relative to the background proteome. Values above zero indicate enrichment, and values below zero depletion. **E** The number of proteins in the SMP set enriched in each amino acid. Insert is a wordcloud summarizing the same data. **F** Density plot showing the distribution of the proportion of sequence length that is low complexity for the SMP proteins (labelled POI for Proteins Of Interest) and the background proteome. **G** Density plot showing the distribution of the proportion of sequence length that is predicted to be intrinsically disordered for the SMP proteins (POI) and the background proteome. **H** The proportion of sequences containing negatively and positively charged clusters of amino acids in the SMP proteins (POI) and the background proteome.

The analysis of sequence bias and amino composition bias with Sequence Properties Analyzer was concordant with the motif finding results, in that G, P and Q were the most enriched amino acids (Figure D). H, I, K and L were found to be the most depleted relative to the background proteome. The most commonly enriched amino acids were P, G, A and C (enriched in 87, 82, 74 and 69 proteins respectively, Figure 2E). Amino acid reidues C and A are not found among the most enriched motifs or the motifs with the highest PS-value, indicating that the proteins are sometimes enriched in an amino acid without that amino acid being embedded in a particular primary sequence context.

The shell matrix proteins showed a clear tendency to contain more low complexity sequence than the background proteome (*p* < 0.0001, Figure 2F) but there was no significant tendency for the sequences to contain a greater proportion of predicted disordered sequence than the background proteome (*p* = 0.1). The shell matrix protein set contained a similar proportion of proteins with negative and positive clusters of amino acids to the background proteome (Figure 2H).

### 3.3 Clustering of proteins based on motif content reveals relationships not found by blast searches

The Sequence Properties Analyser carries out three hierarchical clustering analyses. For one of these, it clusters the proteins based on their enrichment in the 30 most enriched motifs that are present in at least 5% of the proteins. These parameters have been found to yield sensible results across a wide variety of data sets. From the heatmap (Figure 3A) it can immediately be seen that the GG and QP motifs are enriched in a large proportion of the proteins. Despite some noise, 24 clusters were identified in the data (see Materials and Methods). It can be hypothesised that proteins found within the same cluster have a similar function, so we re-ran Protein Motif Finder on each of the clusters in turn to try and extract more biologically meaningful information. The results of five of these analyses are displayed as wordclouds in Figure 3B. Some clusters were dominated by a single motif (e.g. cluster c) while others had much more complex motif profiles (e.g. clusters b and d). Clusters e was particularly rich in M..M motifs, as is immediately obvious from the heatmap. However it was also enriched in SSS..M, T..M, and S.T motifs (among others). None of these motifs were prominent in the analysis of the entire data set, but could be vital to the function of the proteins in this cluster, demonstrating the value of this iterative application of the tools.

**Figure 3:**
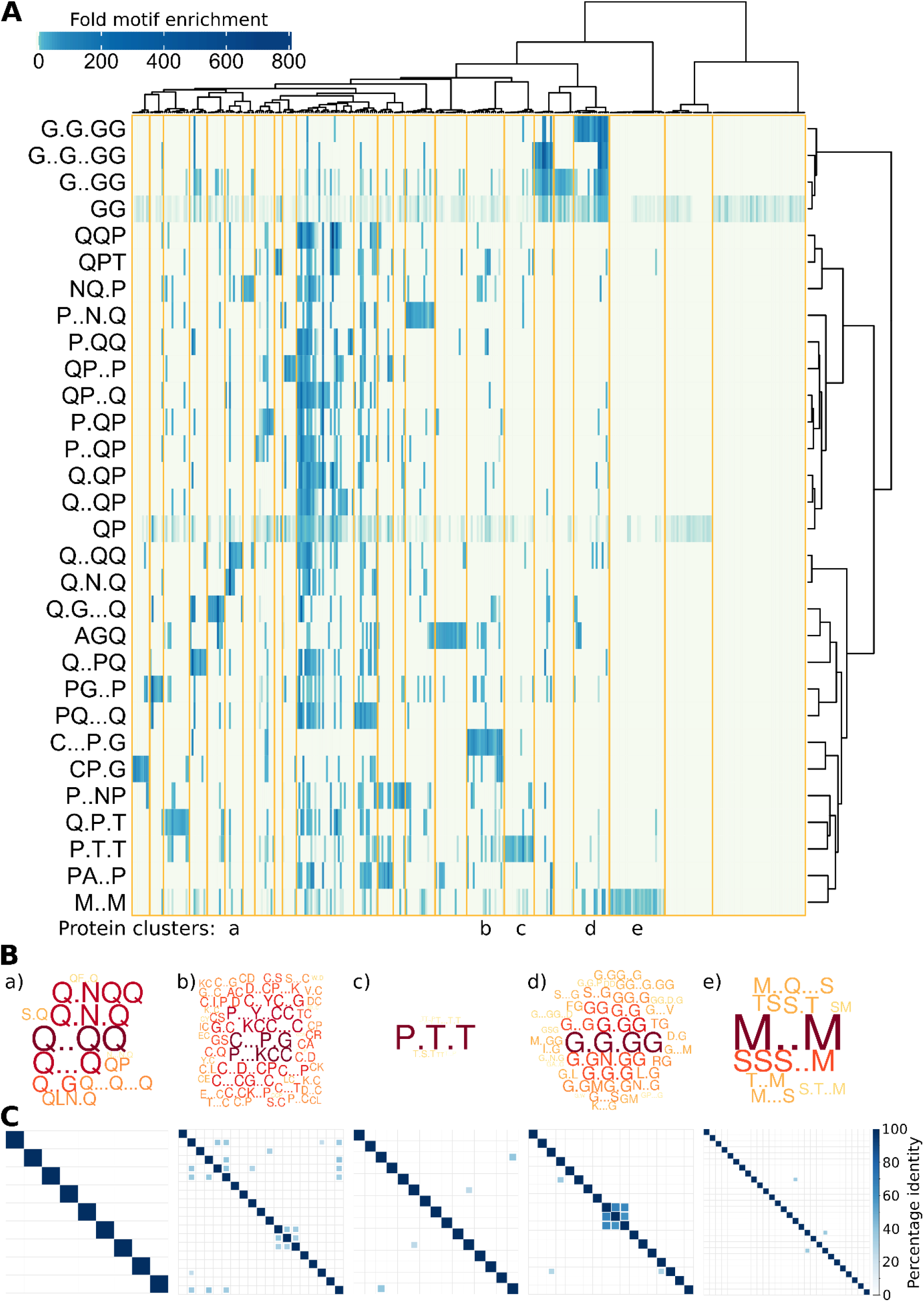
*L. gigantea* shell matrix proteins can be clustered based on motif content despite low sequence identity. The heatmap displays (**A**) motif enrichment in the SMP set relative to the background proteome. Proteins are clustered by their motif enrichment pattern and motifs are clustered by their distribution amongst the proteins. Each motif is a row in the heatmap and each protein is a vertical column. Clusters are marked in gold. For clusters a – e, a wordcloud (**B**) representing the PS-value of the enriched motifs are displayed in addition to a heatmap (**C**) representing the percentage identity between all pairs of proteins in the cluster in an all-vs-all blastp analysis (see Materials and Methods).

We next asked whether the motif clusters reflected larger scale sequence similarity between the proteins within a cluster. To this end, protein sequences in each cluster were subject to an all-vs-all pairwise blastp analysis, which is summarised in the matrices in Figure 3C for five of the clusters. In general larger scale sequence similarity was very low within clusters, with only three protein pairs from the five clusters displaying identity above 50% for the highest scoring HSP.

### 3.4 Detailed analysis of the P.T.T rich cluster using ProminTools

Figure 4 shows the main outputs of Protein Motif Finder for cluster c, which was particularly rich in certain P and T containing motifs. The expected negative correlation between motif enrichment and the number of proteins containing the motif can be seen in figure 4A, for which it is obvious that there is one outlier, P.T.T, which is found in an usually high number of proteins for its enrichment value. We would hypothesise that this motif is particularly important for the function of these proteins, and it is correctly emphasised by the wordcloud based on PS-value in figure 4B. From the heatmap in Figure 4C it is immediately clear that one protein in this cluster is unusual (Lotgi1|227996) in that it has many E containing motifs in addition to the P/T containing motifs. These acidic motifs have much greater fold enrichment than the P/T containing motifs, and so dominate a heatmap based on enrichment (figure 4D), to the extent that other patterns are obscured. Applying a filter requiring that motifs are present in at least 5% of proteins (figure 4B, E) means the significance of the P/T containing motifs can be observed. This analysis illustrates how Protein Motif Finder can facilitate researchers in rapidly detecting the important patterns in their data, including features that are common and features that are unusual within the protein set of interest.

**Figure 4:**
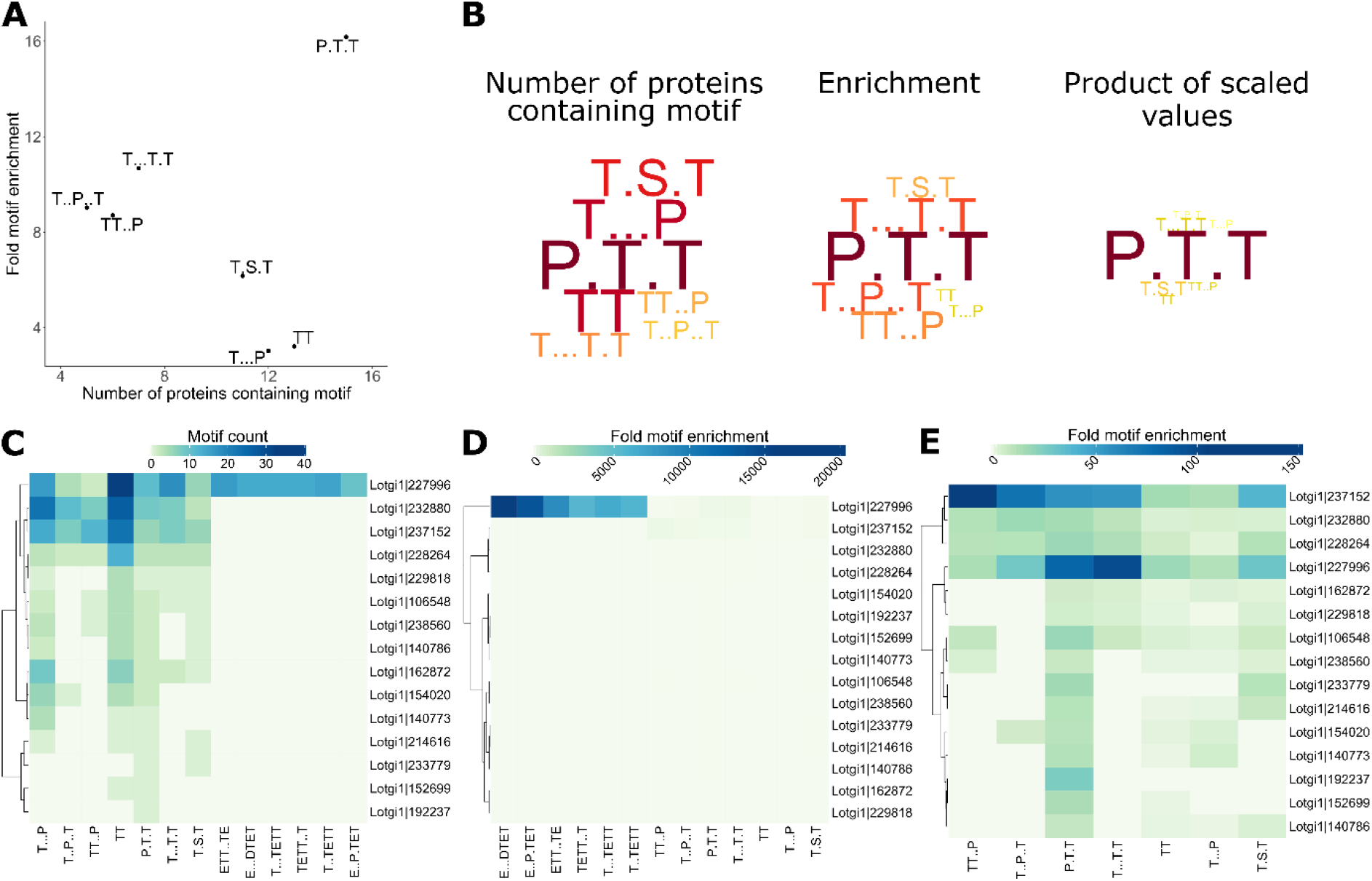
Protein Motif Finder output for cluster c automatically identifies the main features of the data. For cluster c, the relationship between fold motif enrichment and protein count (**A**) and wordclouds visualising these indices (**B**) are displayed. Heatmap **C** displays the count of all enriched motifs in each protein of the cluster; heatmap **D** displays the fold enrichment of all motifs; while heatmap **E** displays the enrichment of only those motifs found in >5% of proteins within the cluster.

## 4. Discussion

Proteins are a prominent part of the organic matrices of many biominerals and are thought to have a number of roles including catalysis, templating, and control of nucleation and crystal growth. Studies of biomineral associated proteins understandably often emphasise proteins with conserved domains, which lend themselves to discussions of their possible molecular functions. However most studies also identify many proteins of unknown function, many of which appear to be low complexity in nature, with biased compositions and a high proportion of intrinsic disorder. Although authors often carefully inspect their protein sequences and note sequences that appear particularly rich in certain residues or motifs, and note the degree of disorder, this information is rarely put in the context of the predicted proteome as a whole.

Here we demonstrate ProminTools, a user friendly package that allows researchers to glean more information from primary sequences of proteins of unknown function and put this in the context of the background proteome. The giant limpet *L. gigantea* has a complex shell matrix proteome for which two different data sets exist. The data analysed in the present study derives from all shell layers (Mann *et al.* [20]), and is thus more complex than the second data set that is derived from the aragonite shell layers only, excluding the calcitic layers (Marie *et al.* [22]). ProminTools revealed a complex array or strongly enriched motifs in the Mann *et al.* data set, which were not uncovered in the original study. Q, P and G rich motifs were particularly prominent and the proteins could be clustered based on their motif content even when they shared little larger scale sequence similarity. Re-running Protein Motif Finder on each of these clusters revealed unique motif profiles that could be hypotheses to be important for the molecular function of proteins in the group. For example one group was enriched in M..M and SSS..M, another in C…P.G and KCC, and another Q..QQ. Interestingly, the Marie *et al*. study also identify a group of low complexity proteins rich in Q and a group rich in M, suggesting that the functions of these proteins may be important for formation of all shell layers or just the aragonite layers, but that they are unlikely to be specific to the calcite layers.

We found that the shell matrix proteins as a group were significantly lower in complexity than the background proteome, providing a statistical underpinning for this observation, and supporting the conclusion of Marie *et al.* who noted the high proportion of low complexity sequences in their data set. The Mann *et al.* studies [12, 20] highlight several proteins in their data which have high degrees of intrinsic disorder. Here, using the Sequence Properties Analyzer we were able to demonstrate that this is not a general feature of the data set, which is not predicted to be significantly more disordered than the background proteome. This highlights the importance of the proteome context when assigning significance to protein features, and demonstrates that the generally observed correlation between protein disorder and low complexity [23] does not hold in every data set.

The role of low complexity regions in biomineralization has only been determined in a very few cases. For example, the enamel protein Amelogenin has a central block of hydrophobic sequence rich in P, H and Q. Intramolecular hydrophobic interactions involving this regions are thought to be critical for self-assembly of Amelogenin into nanospheres and higher order structures that regulated crystal growth [2]. It is possible that the Q and P rich regions in the *L. gigantea* shell matrix proteins might have a similar role in driving self-assembly processes.

Although at present we can only speculate on the role of low complexity proteins in biomineralization, it is clear that low complexity sequences are not unique to biomineralization related proteins. Depending on the species, 22 – 36 % of residues in eukaryotic proteins fall into low complexity regions [24]. It remains to be investigated whether the low complexity regions of biomineralization related proteins have features that set them apart from other low complexity regions in proteomes, and ProminTools could be used to investigate such questions.

ProminTools allows researchers to easily find patterns in their data, but it has limitations and researchers should use their judgement in interpreting the output. For example the hierarchical clustering method will always generate some clusters whether or not there are interesting patterns in the data, and users must inspect the output to decide if the results are meaningful. It should also be remembered that this is a tool for hypothesis generation. For example, proteins which share similar motifs can be hypothesised to perform similar molecular functions, but this may or may not be the case in reality and experimental validation is required. ProminTools will be at its most useful when combined with other methods for spotting repeating patterns in sequences (eg. HhpreID [25], Meme [26] or simply inspecting dot-plots) and when put in the context of additional information such as known domain content, post-translational modifications, phylogenetic distributions and expression patterns.

We would like to point out that ProminTools can be used for any pairwise comparison of sets of protein sequences. For examples, protein sets associated with different part of a biomineral or different developmental stages could also be compared, and if carefully carried out, cross-species comparisons could also be made. The latter could be particularly useful, since the fast evolving nature of low complexity sequences [27] can make it difficult to detect homology. It could also be applied to other protein sets rich in low complexity sequences, such as proteins found in pathological amyloids associated with diseases such as Alzheimer’s and Parkinson’s [28].

In conclusion, it is clear that ProminTools will help researchers generate new hypotheses about the important of particular motifs and protein chemistries in their system of interest and provide new directions for experimental work. Putting the patterns identified into the context of the rest of the proteome ensures that features that are genuinely overrepresented in the POIs are prioritised for further study.

## Acknowledgements

We gratefully acknowledge Dr. A. Scheffel for advice and critical reading of the manuscript. AS acknowledges financial support from the Alexander von Humboldt foundation.

## Author contributions

AS conceived the study, wrote the tools, compiled the figures and wrote the manuscript. AD containerised the tools and reviewed the manuscript.

## Software availability

ProminTools is available in Apps with graphical user interfaces from the Cyverse Discovery Environment (https://de.cyverse.org/). Users need to create an account, but then have access to a large number of tools and high performance computing. Docker images are available from Docker Hub (https://hub.docker.com/repositories/biologistatsea/) and detailed instructions on usage and uses are available at https://github.com/skeffington/Promin-tools.

## Supporting Information

Supplementary file 1 contains the fasta file of the *L. gigantea* SMP used in this study: Supp1_Lg_matrix_proteins.fasta

Supplementary file 2 contains the full Protein Motif Finder output for the *L. gigantea* SMPs: Supp2_LgMP_motif_finder.zip

Supplementary file 3 contains the full Sequence Properties Analyzer output for the *L. gigantea* SMPs: Supp3_LgMP_seq_prop.zip

Supplementary file 4 contains the Protein Motif Finder output for all clusters: Supp4_LgMP_clusters_motif_finder.zip

Supplementary file 5 contains the licence information for software used by ProminTools: Supp5_licence_information.txt

## List of Abbreviations

SVG: Scalable Vector Graphic
POI: Protein Of Interest
SMP: Shell Matrix Protein
HSP: High Scoring Pair

