## Supplemental File 3 for "ProminTools: Shedding light on proteins of unknown function in biomineralization with user friendly tools illustrated using mollusc shell matrix protein sequences": Mann2014all_Rscript.R

```
#' ---
#' title: Properties of your proteins of interest relative to background sequences
#' author: A tool by Alastair Skeffington hosted by Cyverse
#' output: 
#'  html_document:
#'    toc: TRUE
#'    toc_float: TRUE
#' ---
#'
#'
#'<style>
#'  body {
#'    text-align: justify}
#'</style>
#'
#'
#'<style>
#'  p.caption {
#'  font-size: 0.9em;
#'  font-style: italic;
#'  color: grey;
#'  margin-right: 10%;
#'  margin-left: 10%;  
#'  text-align: justify;
#'  }
#'</style>


#+ include = FALSE
library (wordcloud)
library(dplyr)
library(ggplot2)
library(DT)
library(forcats)
library(kableExtra)
library(ggrepel)
library(gplots)
library(knitr)

library(ComplexHeatmap)


#'
#'## Introduction


#'This tool aims to generate useful statistical information about a set of protein sequences of interest (POIs)
#'with respect to a background set of proteins, for example the predicted proteome of the species concerned.
#'  
#'<br>
#'The main tabular outputs of the program are listed below:  
#+ echo = FALSE, warnings=FALSE, messages=FALSE

#'* Mann2014all _AAabundance.txt: The data displayed in table 2.
#'* Mann2014all _AAenrich.txt: Enrichment (fold change) for each amino acid in the POI with respect to the background (displayed in table 1). The second numeric column is the data for figure 1.
#'* Mann2014all _bg.comp: Relative frequencies of amino acids in the background protein set
#'* Mann2014all _fg.comp: Relative frequencies of amino acids in the POI protein set
#'* Mann2014all _chargedclus_POI_summary.txt: For each protein in the POI set, the number, length and percentage of the sequence covered by positive and negative clusters of amino acids. Data used in figure 5.
#'* Mann2014all _chargedclus_POI.txt: The list of POI proteins containing positive or negative clusters of amino acids, along with the positions of the clusters.
#'* Mann2014all _chargedclus_Proteome_summary.txt: For each protein in the background proteome set, the number, length and percentage of the sequence covered by positive and negative clusters of amino acids. Data used in figure 5.
#'* Mann2014all _chargedclus_Proteome.txt: The list of background proteome proteins containing positive or negative clusters of amino acids, along with the positions of the clusters.
#'* Mann2014all _chargedclus_seqtbl.txt: The data displayed in table 4.
#'* Mann2014all _complexityPOI_summary.txt: The data displayed in table 3. Data used in figure 3.
#'* Mann2014all _complexityProteome_summary.txt: Similar to the data displayed in table 3, but for the background proteome protein set. Data used in figure 3.
#'* Mann2014all _POI_disorder_summary.txt: For each POI protein, gives the length and number of disordered regions and the percentage of the sequence predicted to be disordered.
#'* Mann2014all _POI_disorder.txt: Gives the position in the POI proteins of each disordered region. Data used in figure 4.
#'* Mann2014all _Proteome_disorder_summary.txt: For each background proteome protein, gives the length and number of disordered regions and the percentage of the sequence predicted to be disordered.
#'* Mann2014all _Proteome_disorder.txt: Gives the position in the background proteome proteins of each disordered region. Data used in figure 4.
#'* Mann2014all _positions.txt: Gives the positions of all compositionally biased regions in the POI protein set, including the residues that are biased and the inverse of the binomial p-value for the bias.
#'* Mann2014all _protcount.txt: For each type of compositional bias (amino acid or group of amino acids), gives the count of proteins in the POI set containing this bias. Data used in figure 2.
#'* Mann2014all _Rscript.R : The R script that generated the html output. This can be used as a basis to produce your own custom figures and plots in R  

#+ echo = FALSE, warnings=FALSE, messages=FALSE

#'
#'<br>
#'The main outputs of the program are the data tables. For convenience these are processed to an html document with
#'some visualizations of the data. However one analysis pipeline can never be appropriate for every dataset,
#'so the R script used to generate these diagrams is also an output of the program. This is to allow you, or an
#'infromatically minded colleague to easily tweak the plots or to develop these analyses further in a way 
#'appropriate for your data.  
#'<br>
#'
#'
#+ echo = FALSE, warnings=FALSE, messages=FALSE
AAenrich.path = "/home/mpimp-golm.mpg.de/skeffington/winhome/diatoms/cyverse_tools/seq_properties2/Mann2014all_AAenrich.txt"
AAprotcount.path = "/home/mpimp-golm.mpg.de/skeffington/winhome/diatoms/cyverse_tools/seq_properties2/Mann2014all_protcount.txt"
AAabun.path = "/home/mpimp-golm.mpg.de/skeffington/winhome/diatoms/cyverse_tools/seq_properties2/Mann2014all_AAabundance.txt"
Bias_index.path = "/home/mpimp-golm.mpg.de/skeffington/winhome/diatoms/cyverse_tools/seq_properties2/Mann2014all_biasindex.txt"
Compl_poi.path = "/home/mpimp-golm.mpg.de/skeffington/winhome/diatoms/cyverse_tools/seq_properties2/Mann2014all_complexityPOI_summary.txt"
Compl_prot.path = "/home/mpimp-golm.mpg.de/skeffington/winhome/diatoms/cyverse_tools/seq_properties2/Mann2014all_complexityProteome_summary.txt"
dis_poi.path = "/home/mpimp-golm.mpg.de/skeffington/winhome/diatoms/cyverse_tools/seq_properties2/Mann2014all_POI_disorder_summary.txt"
dis_prot.path = "/home/mpimp-golm.mpg.de/skeffington/winhome/diatoms/cyverse_tools/seq_properties2/Mann2014all_Proteome_disorder_summary.txt"
chcl_poi.path = "/home/mpimp-golm.mpg.de/skeffington/winhome/diatoms/cyverse_tools/seq_properties2/Mann2014all_chargedclus_POI_summary.txt"
chcl_prot.path = "/home/mpimp-golm.mpg.de/skeffington/winhome/diatoms/cyverse_tools/seq_properties2/Mann2014all_chargedclus_Proteome_summary.txt"
chcl_seqble.path = "/home/mpimp-golm.mpg.de/skeffington/winhome/diatoms/cyverse_tools/seq_properties2/Mann2014all_chargedclus_seqtbl.txt"


#+ echo = FALSE, warnings=FALSE, messages=FALSE
#'## Compositional bias
#'The following graph displays the degree of enrichment of each amino acid in the POI set verses the 
#'background protein set. Values above zero indicate relative enrichment which bars below zero indicate
#'relative depletion. Absolute enrichment values (fold change in frequency) are given in the table.
#'  
#'
#'
#+ echo = FALSE, warnings=FALSE, messages=FALSE, fig.cap="Figure 1: Bias in amino acid composition"
AAenrich<-read.table(AAenrich.path, sep="\t", header=FALSE)
AAbiasplot<-ggplot(AAenrich, aes(V1, V3))
AAbiasplot + geom_bar(width=0.8, position = "dodge", stat="identity", fill= "#34b4eb") + 
  theme_classic() +
  ylab("(POI set frequency - background frequency) /\n background frequency") + 
  xlab("Amino acid") + geom_hline(yintercept = 0) 

AAenrich<-AAenrich[order(AAenrich$V2, decreasing = TRUE), ]
AAenrich$Enrichment<-round(AAenrich$V2, 2)
AAenrich_2<-select(AAenrich, V1, Enrichment)
colnames(AAenrich_2)<-c("Amino acid", "Enrichment")

#'
#'
#'<br>
#'<br>
#+ echo = FALSE, warnings=FALSE, messages=FALSE, fig.cap="Table 1: Bias in amino acid composition"
kable(AAenrich_2, row.names=FALSE, colnames=c("Amino acid", "Fold enrichment")) %>%
    kable_styling(bootstrap_options = c("striped", "hover", "condensed"), full_width = F) %>%
    footnote(general = "", general_title = "Table 1: Bias in amino acid composition",
           footnote_as_chunk = T, title_format = c("italic"))

#'Below, the number of proteins enriched in each amino acid (or group of amino acids) is displayed as
#'a wordcloud (left, letter height relates to protein number) or a bar graph (right). In the bar graph,
#'only the 20 amino acids enriched in the greatest number of proteins are displayed. In the wordcloud a 
#'residue or group of residues must be present in at least in 10% of the proteins in order to be displayed.
 
#+ echo = FALSE, warnings=FALSE, messages=FALSE 
AAprotcount<-read.table(AAprotcount.path, sep="\t", header=TRUE)
colnames(AAprotcount)<-c("Residues", "Pcount")

if( length(AAprotcount$Residues) < 20){
  AAprotcount_grmax = length(AAprotcount$Residues)
}else{
  AAprotcount_grmax = 20
}

#+echo = FALSE, warnings=FALSE, messages=FALSE
AAprotcount2<-AAprotcount[order(AAprotcount$Pcount, decreasing = TRUE),]
AAprotcount2<-head(AAprotcount2, AAprotcount_grmax)
AAprotcount2$Residues <- factor(AAprotcount2$Residues, levels = AAprotcount2$Residues[order(AAprotcount2$Pcount, decreasing = TRUE)])
AAprotcount2_plot<-ggplot(AAprotcount2, aes(Residues, Pcount)) + 
  theme_classic() +
  geom_bar(width=0.8, position = "dodge", stat="identity", fill= "#34b4eb") +
  ylab("Number of protein enriched") +
  xlab("")


#+r, out.width=c('40%', '60%'), fig.show='hold', echo = FALSE, warnings=FALSE, messages=FALSE, fig.cap="Figure 2: Proportion of proteins displaying bias in given amino acids or groups of amino acids"
wordcloud(AAprotcount$Residues, AAprotcount$Pcount, random.order=FALSE, scale = c(10, 0.001),
          rot.per="0", colors= brewer.pal(12, "Paired"), min.freq = 0.2)
AAprotcount2_plot
#'<br>
#'
#'
#+ echo = FALSE, warnings=FALSE, messages=FALSE
AAabundtbl<-read.table(AAabun.path, sep="\t", header=TRUE)


#'<br>
#'The interactive table below displays the top 3 most abundant residues in each protein (abundance rank 1 - 3) along with
#'their percentage abundance. It also displays the 3 most enriched residues for each protein, along with
#'the fold change in frequency relative to the background proteome (Enrichment_FC_) and also their percentage
#'abundance in the protein sequence (En_percentage_abun_).
#'<br>
#+ echo = FALSE, warnings=FALSE, messages=FALSE, fig.cap="Table 2: Bias in amino acid composition for each protein"
round_df <- function(df, digits) {
  nums <- vapply(df, is.numeric, FUN.VALUE = logical(1))
  
  df[,nums] <- round(df[,nums], digits = digits)
  
  (df)
}
datatable(round_df(AAabundtbl, 2), rownames = TRUE, filter="top", options = list(pageLength = 8, scrollX=T))

#'<br>
#+ echo = FALSE, warnings=FALSE, messages=FALSE
biasindex<-read.table(Bias_index.path, sep="\t", header=TRUE)
biasden <- density.default(biasindex[biasindex$Source == "background", ]$bias_index, n = 512, cut = 3)
xbiasden <- biasden$x  
ybiaseden <- biasden$y 

fgbiasindex<-biasindex[biasindex$Source == "forground", ]$bias_index
fgbiasindex<-round(fgbiasindex, digits = 2)
fbias <- approxfun(xbiasden, ybiaseden)
Cbias <- integrate(fbias, min(xbiasden), max(xbiasden))$value
if(fgbiasindex < max(xbiasden)){
  pbi.unscaled <- integrate(fbias, 0.16, max(xbiasden))$value
  pbi.scaled <- pbi.unscaled / Cbias
}else{
  pbi.scaled <- 0
}
pbi.scaled<-round(pbi.scaled, digits = 4)
if(pbi.scaled == 0){
	pbi.scaled <- "<0.0001"
}

#'<br>
#'A measure of the degree of bias is given by the bias index, defined as the square root of the sum (across all amino acids) of squared deviations
#'in amino acid frequency between the POI set and the background proteome. For this data, the bias index is `r fgbiasindex`. A bias index of zero would indicate
#'no deviation from the background amino acid frequencies. By repeated sampling of the background proteome and calculating the bias index for 
#'these samples, the frequency distribution of bias index values you would expect to occur by random sampling of the proteome can be approximated,
#'allowing a p-value for the degree of bias in the POI set to be estimated. This p-value is the probability of getting a protein set as biased as or
#'more extreme than the POI set via random sampling of the background proteome. The p-value for this POI set is: `r pbi.scaled`.
#'

#'<br>
#'
#'
#'## Sequence complexity
#'
#'Sequence complexity was analysed using the program [Seg](http://www.biology.wustl.edu/gcg/seg.html). 
#'Sequences below a certain complexity threshold were extracted. The density plot displays the distribution
#'of lengths of low complexity sequence among proteins from the set of proteins of interest (POIs) and the
#'background sequence set.
#+echo = FALSE, warnings=FALSE, messages=FALSE,fig.cap="Figure 3: Density plots of the percentage of low complexity sequence"
Compl_poi<-read.table(Compl_poi.path, sep="\t", header=TRUE)
Compl_prot<-read.table(Compl_prot.path, sep="\t", header=TRUE)

Compl_poi_lcrpc<-select(Compl_poi, lcr_percnt)
Compl_prot_lcrpc<-select(Compl_prot, lcr_percnt)
Compl_poi_lcrpc$Source<-rep("POIs")
Compl_prot_lcrpc$Source<-rep("Background sequences")
compl_data<-rbind(Compl_poi_lcrpc, Compl_prot_lcrpc)
Segdensity<-ggplot()
Segdensity + geom_density(data=compl_data, aes(x = lcr_percnt, y = ..scaled.., fill = fct_rev(Source)), alpha = 0.5) + theme_classic() + scale_y_continuous(expand = c(0,0)) + 
  scale_x_continuous(expand = c(0,0)) +
  xlab("Percentage of sequence which is low complexity") + ylab("Scaled density") + labs(fill = "Source")
#+echo = FALSE, warnings=FALSE, messages=FALSE
compl_wilk<-wilcox.test(Compl_poi_lcrpc$lcr_percnt, Compl_prot_lcrpc$lcr_percnt)
compl_wilk_pval<-round(compl_wilk$p.value, 4)
if(compl_wilk_pval == 0){
	compl_wilk_pval <- "<0.0001"
}

#'We can use a Wilcoxon rank sum test with continuity correction to test if the difference in complexity 
#'between the POIs and the background sequences is significant. The null hypothesis is that there is no 
#'difference in complexity betweeen the two sets of proteins. The p value respresents the probability of 
#'observing the data if the null hypothesis is true, so a low p-value allows us to reject the null hypothesis.
#'The pvalue is this case is  `r compl_wilk_pval`.
#'<br>
#'
#'The table below present, for each protein, the total length of the low complexity regions in the protein,
#'the percentage of the sequence length that is low complexity, the average complexity of the low complexity 
#'sequence (measured in bits) and the enrichment (fold change in density) of each amino acid in the low 
#'complexity sequence realtive to the background proteome. The table is interactive and should allow easy
#'exploration of the data.
#'<br>
#'
#'
#+echo = FALSE, warnings=FALSE, messages=FALSE, fig.cap="Table 3: Low complexity regions in the POI sequence set"
rownames(Compl_poi) <- Compl_poi[,1]
Compl_poi <- Compl_poi %>% 
            rename(
              LCR_length = lcr_len,
              LCR_percent = lcr_percnt,
              LCR_complexity = lcr_complexity
  )
Compl_poi[,1]<-NULL
datatable(round(Compl_poi, 2), rownames = TRUE, filter="top", options = list(pageLength = 8, scrollX=T))


#'## Predicted intrinsic disorder
#'The intrinsic disorder of proteins was assessed using the [VSL2](https://bmcbioinformatics.biomedcentral.com/articles/10.1186/1471-2105-7-208) predictor
#'of the [DisProt](http://www.dabi.temple.edu/disprot/predictor.php) family of predictors. Although there are
#'more modern methods available, this method has good accuracy and the advantage of being very fast, making
#'the calculation of disorder across a whole proteome feasible in a reasonable length of time. The density plot 
#'displays the distribution of the percentage of the sequence that is disordered among proteins from the 
#'set of proteins of interest (POIs) and the background sequence set.
#'
#+echo = FALSE, warnings=FALSE, messages=FALSE, fig.cap="Figure 4: Density plots of the percent of disordered sequence"
dis_poi<-read.table(dis_poi.path, sep = "\t", header=TRUE)
dis_prot<-read.table(dis_prot.path, sep = "\t", header=TRUE)

dis_poi$Source<-rep("POIs")
dis_prot$Source<-rep("Proteome")
dis_all<-bind_rows(select(dis_poi, disordered_percent, Source), select(dis_prot, disordered_percent, Source))

vsl2plot<-ggplot()
vsl2plot + geom_density(data=dis_all, aes(x = disordered_percent, y = ..scaled.., fill = Source), alpha = 0.5) + theme_classic() +
   scale_y_continuous(expand = c(0,0)) + scale_x_continuous(expand = c(0,0)) +
   xlab("Percentage of sequence which is disordered") + ylab("Scaled density")

dis_wilk<-wilcox.test(dis_poi$disordered_percent, dis_prot$disordered_percent)
dis_wilk_pval<-round(dis_wilk$p.value, 4)
if(dis_wilk_pval == 0){
	dis_wilk_pval <- "<0.0001"
}

#'We can use a Wilcoxon rank sum test with continuity correction to test if the difference in disorder 
#'between the POIs and the background sequences is significant. The null hypothesis is that there is no 
#'difference in disorder between the two sets of proteins. The p-value represents the probability of 
#'observing the data if the null hypothesis is true, so a low p-value allows us to reject the null hypothesis.
#'The pvalue is this case is  `r dis_wilk_pval`.
#'<br>
#'

#'## Charged clusters
#'Clusters of charged amino acids were identified using [SAPS](https://www.pnas.org/content/89/6/2002). The
#'graph below shows the percentage of sequences in the POI set and the background proteome that contain
#'charged clusters.
#'<br>
#+echo = FALSE, warnings=FALSE, messages=FALSE, fig.cap="Figure 5: Percentage of proteins with charged clusters"
chcl_poi<-read.table(chcl_poi.path, sep = "\t", header=TRUE)
chcl_prot<-read.table(chcl_prot.path, sep = "\t", header=TRUE)

chcl_neg <- matrix(c(sum((chcl_poi$number_neg_cluster > 0), na.rm = TRUE), sum((chcl_prot$number_neg_cluster > 0), na.rm = TRUE), 
                     length(unique(chcl_poi$name)), length(unique(chcl_prot$name))), ncol = 2)

chcl_neg<-as.data.frame(chcl_neg)
chcl_neg$pc<-chcl_neg$V1 / chcl_neg$V2 *100
chcl_neg$charge<-rep("negative")

chcl_pos <- matrix(c(sum((chcl_poi$number_pos_cluster > 0), na.rm = TRUE), sum((chcl_prot$number_pos_cluster > 0), na.rm = TRUE), 
                     length(unique(chcl_poi$name)), length(unique(chcl_prot$name))), ncol = 2)
chcl_pos<-as.data.frame(chcl_pos)
chcl_pos$pc<-chcl_pos$V1 / chcl_pos$V2 *100
chcl_pos$charge<-rep("positive")

chcl_all<-rbind(chcl_neg, chcl_pos)
chcl_all$Source<-c("POI", "Proteome", "POI", "Proteome")

chcl_plot<-ggplot(chcl_all, aes(charge, pc))
chcl_plot + geom_bar(aes(fill = Source), position="Dodge", stat="identity") + theme_classic() +
  scale_y_continuous(expand = c(0,0)) +
  ylab("Percentage of sequences\nconataining charged cluster") + xlab("Charge")

#'
#'<br>
#'
#'Those proteins that contain charged clusters, along with the sequence of the cluster and the position of
#'the sequence in the protein, are displayed in the table below.
#'<br>
#'
#+echo = FALSE, warnings=FALSE, messages=FALSE, fig.cap="Table 4: Charged clusters in the POI sequences"
chcl_seqble<-read.table(chcl_seqble.path, sep = "\t", header=TRUE)
datatable(chcl_seqble, rownames = TRUE, filter="top", options = list(pageLength = 8, scrollX=T))
```
