## Supplemental File 3 for "ProminTools: Shedding light on proteins of unknown function in biomineralization with user friendly tools illustrated using mollusc shell matrix protein sequences": Mann2014all_sequence_properties.html

Properties of your proteins of interest relative to background sequences


### Properties of your proteins of interest relative to background sequences

###### A tool by Alastair Skeffington hosted by Cyverse

#### 2020-02-26

#### Introduction

This tool aims to generate useful statistical information about a set of protein sequences of interest (POIs) with respect to a background set of proteins, for example the predicted proteome of the species concerned.

The main tabular outputs of the program are listed below:

- Mann2014all \_AAabundance.txt: The data displayed in table 2.
- Mann2014all \_AAenrich.txt: Enrichment (fold change) for each amino acid in the POI with respect to the background (displayed in table 1). The second numeric column is the data for figure 1.
- Mann2014all \_bg.comp: Relative frequencies of amino acids in the background protein set
- Mann2014all \_fg.comp: Relative frequencies of amino acids in the POI protein set
- Mann2014all \_chargedclus\_POI\_summary.txt: For each protein in the POI set, the number, length and percentage of the sequence covered by positive and negative clusters of amino acids. Data used in figure 5.
- Mann2014all \_chargedclus\_POI.txt: The list of POI proteins containing positive or negative clusters of amino acids, along with the positions of the clusters.
- Mann2014all \_chargedclus\_Proteome\_summary.txt: For each protein in the background proteome set, the number, length and percentage of the sequence covered by positive and negative clusters of amino acids. Data used in figure 5.
- Mann2014all \_chargedclus\_Proteome.txt: The list of background proteome proteins containing positive or negative clusters of amino acids, along with the positions of the clusters.
- Mann2014all \_chargedclus\_seqtbl.txt: The data displayed in table 4.
- Mann2014all \_complexityPOI\_summary.txt: The data displayed in table 3. Data used in figure 3.
- Mann2014all \_complexityProteome\_summary.txt: Similar to the data displayed in table 3, but for the background proteome protein set. Data used in figure 3.
- Mann2014all \_POI\_disorder\_summary.txt: For each POI protein, gives the length and number of disordered regions and the percentage of the sequence predicted to be disordered.
- Mann2014all \_POI\_disorder.txt: Gives the position in the POI proteins of each disordered region. Data used in figure 4.
- Mann2014all \_Proteome\_disorder\_summary.txt: For each background proteome protein, gives the length and number of disordered regions and the percentage of the sequence predicted to be disordered.
- Mann2014all \_Proteome\_disorder.txt: Gives the position in the background proteome proteins of each disordered region. Data used in figure 4.
- Mann2014all \_positions.txt: Gives the positions of all compositionally biased regions in the POI protein set, including the residues that are biased and the inverse of the binomial p-value for the bias.
- Mann2014all \_protcount.txt: For each type of compositional bias (amino acid or group of amino acids), gives the count of proteins in the POI set containing this bias. Data used in figure 2.
- Mann2014all \_Rscript.R : The R script that generated the html output. This can be used as a basis to produce your own custom figures and plots in R

The main outputs of the program are the data tables. For convenience these are processed to an html document with some visualizations of the data. However one analysis pipeline can never be appropriate for every dataset, so the R script used to generate these diagrams is also an output of the program. This is to allow you, or an infromatically minded colleague to easily tweak the plots or to develop these analyses further in a way appropriate for your data.

#### Compositional bias

The following graph displays the degree of enrichment of each amino acid in the POI set verses the background protein set. Values above zero indicate relative enrichment which bars below zero indicate relative depletion. Absolute enrichment values (fold change in frequency) are given in the table.

Figure 1: Bias in amino acid composition

| Amino acid | Enrichment |
| --- | --- |
| P | 1.31 |
| G | 1.27 |
| Q | 1.19 |
| A | 1.17 |
| C | 1.12 |
| T | 1.08 |
| D | 1.04 |
| N | 1.04 |
| V | 1.03 |
| M | 1.02 |
| Y | 0.98 |
| S | 0.96 |
| R | 0.95 |
| F | 0.92 |
| E | 0.89 |
| W | 0.88 |
| I | 0.86 |
| K | 0.84 |
| L | 0.82 |
| H | 0.73 |
|  |  |
| --- | --- |
| Table 1: Bias in amino acid composition |  |

Below, the number of proteins enriched in each amino acid (or group of amino acids) is displayed as a wordcloud (left, letter height relates to protein number) or a bar graph (right). In the bar graph, only the 20 amino acids enriched in the greatest number of proteins are displayed. In the wordcloud a residue or group of residues must be present in at least in 10% of the proteins in order to be displayed.

Figure 2: Proportion of proteins displaying bias in given amino acids or groups of amino acids

The interactive table below displays the top 3 most abundant residues in each protein (abundance rank 1 - 3) along with their percentage abundance. It also displays the 3 most enriched residues for each protein, along with the fold change in frequency relative to the background proteome (Enrichment\_FC\_) and also their percentage abundance in the protein sequence (En\_percentage\_abun\_).

Table 2: Bias in amino acid composition for each protein

A measure of the degree of bias is given by the bias index, defined as the square root of the sum (across all amino acids) of squared deviations in amino acid frequency between the POI set and the background proteome. For this data, the bias index is 0.12. A bias index of zero would indicate no deviation from the background amino acid frequencies. By repeated sampling of the background proteome and calculating the bias index for these samples, the frequency distribution of bias index values you would expect to occur by random sampling of the proteome can be approximated, allowing a p-value for the degree of bias in the POI set to be estimated. This p-value is the probability of getting a protein set as biased as or more extreme than the POI set via random sampling of the background proteome. The p-value for this POI set is: <0.0001.

#### Sequence complexity

Sequence complexity was analysed using the program Seg. Sequences below a certain complexity threshold were extracted. The density plot displays the distribution of lengths of low complexity sequence among proteins from the set of proteins of interest (POIs) and the background sequence set.

Figure 3: Density plots of the percentage of low complexity sequence

We can use a Wilcoxon rank sum test with continuity correction to test if the difference in complexity between the POIs and the background sequences is significant. The null hypothesis is that there is no difference in complexity betweeen the two sets of proteins. The p value respresents the probability of observing the data if the null hypothesis is true, so a low p-value allows us to reject the null hypothesis. The pvalue is this case is <0.0001.

The table below present, for each protein, the total length of the low complexity regions in the protein, the percentage of the sequence length that is low complexity, the average complexity of the low complexity sequence (measured in bits) and the enrichment (fold change in density) of each amino acid in the low complexity sequence realtive to the background proteome. The table is interactive and should allow easy exploration of the data.

Table 3: Low complexity regions in the POI sequence set

#### Predicted intrinsic disorder

The intrinsic disorder of proteins was assessed using the VSL2 predictor of the DisProt family of predictors. Although there are more modern methods available, this method has good accuracy and the advantage of being very fast, making the calculation of disorder across a whole proteome feasible in a reasonable length of time. The density plot displays the distribution of the percentage of the sequence that is disordered among proteins from the set of proteins of interest (POIs) and the background sequence set.

Figure 4: Density plots of the percent of disordered sequence

We can use a Wilcoxon rank sum test with continuity correction to test if the difference in disorder between the POIs and the background sequences is significant. The null hypothesis is that there is no difference in disorder between the two sets of proteins. The p-value represents the probability of observing the data if the null hypothesis is true, so a low p-value allows us to reject the null hypothesis. The pvalue is this case is 0.1144.

#### Charged clusters

Clusters of charged amino acids were identified using SAPS. The graph below shows the percentage of sequences in the POI set and the background proteome that contain charged clusters.

Figure 5: Percentage of proteins with charged clusters

Those proteins that contain charged clusters, along with the sequence of the cluster and the position of the sequence in the protein, are displayed in the table below.

Table 4: Charged clusters in the POI sequences
