## Supplemental File 4 for "ProminTools: Shedding light on proteins of unknown function in biomineralization with user friendly tools illustrated using mollusc shell matrix protein sequences": Mann14_cl1_motifs.html

Overrepresented motifs in your proteins of interest


### Overrepresented motifs in your proteins of interest

###### A tool by Alastair Skeffington hosted by Cyverse and making use of the motif-x motif finding engine

#### 2020-03-03

The main outputs of the program are listed below:

- Mann14\_cl1 \_bgmotifs.txt : counts of each of the overrepresented motifs in each of the proteins in the background sequence set
- Mann14\_cl1 \_fgmotifs.txt : counts of each of the overrepresented motifs in each of the proteins in the foreground sequence set
- Mann14\_cl1 \_fgenrich.txt : the enrichment of each of the overrepresented motifs in each of the proteins in the foreground sequence set with repspect to the background sequence set
- Mann14\_cl1 \_motifsummary.txt : for each overrepresented motif, the enrichemnt in the forgraound sequnces relative to the background; the count of proteins in which it appears and the median count of the motif per protein
- Mann14\_cl1 \_motifs.html : A html summary of the analysis with some helpful plots
- Mann14\_cl1 \_Rscript.R : The R script that generated the html output. This can be used as a basis to produce your own custom figures and plots in R
- Mann14\_cl1 \_wordclouds.svg : An svg file of the wordclouds from the html report. This can be imported into Inkscape or other software to make publication ready figures

The main outputs of the program are the data tables. For convenience these are processed to a html document with some visualizations of the data. However one analysis pipeline can never be appropriate for every dataset, so the R script used to generate these diagrams is also an output of the program. This is to allow you, or an infromatically minded colleague to easily tweak the plots or to develop these analyses further in a way appropriate for your data.

| Motif | Enrichment | Count of proteins with motif | Median motif count per protein |
| --- | --- | --- | --- |
| R..G | 2.24 | 32 | 1 |
| S…G | 1.80 | 35 | 1 |
| G…S | 1.51 | 35 | 1 |

The data in the table above can be visualized as wordclouds. To avoid plotting problems, any infinitely enriched motifs were first removed from the data. In this analyses we are not interested in the peculiarities of particular proteins, but in the properties of this group of proteins as a whole, so any motifs found in less than 5% of the proteins or in only one protein, were also removed before plotting.

In the wordcoulds the height of the letters relates to either the number of proteins in the set of interest containing a given motif (left), the enrichment of a motif relative to the background proteome (middle), or the product of the scaled values of these two measures (right). In some datasets with very few motifs, the scaled values are numerically undefined and no wordcloud will be plotted.

In general there tends to be a negative correlation between the enrichment of a motif and the number of proteins in which it is found. It is possible that motifs that deviate from this trend (ie they are unusually enriched given the number of proteins in which they are found, or are in an unusually large number of proteins given their enrichment) might have particular biological significance. The scatter plots below plot the enrichment of the motifs against the number of proteins in which they are found (and vice versa) to help you visualize any deviations from the expected negative correlation. If a linear regression is appropriate, a regression line (either linear or polynomial) is shown in red and the shaded area represents the 95% confidence interval of the regression. No regression line is shown in cases with strong non-linearity. Points are labelled only if there is sufficient space in the plot.

You may be interested in grouping or subdividing your PSOI based on the motif content of the proteins. The heatmaps below provide a starting point for such approaches. The top heatmap displays a hierarchical clustering of the proteins based on the number of motifs they contain, and simultaneously a clustering of motifs based on their distributions amongst proteins. In contrast to the above analyses, the filters that remove the infinitely enriched motifs and motifs found in less than 5% of proteins are not applied. For clarity however, only the 30 motifs found most frequently amongst the proteins are used in the clustering for the first heatmap. Proteins containing none of the motifs are not displayed.

In the second heatmap, proteins are clustered based on motif enrichment in a given protein sequence with respect the the background sequences, and motifs are clustered based on their enrichment amongst proteins. The filter that removes motifs found in less than 5% of proteins is not applied, but infinitely enriched motifs are excluded. However clustering is restricted to the 30 motifs with the greatest overall enrichment in the dataset. Proteins containing none of the motifs are not displayed.

In the third heatmap, proteins are clustered based on motif enrichment in a given protein sequence with respect the the background sequences, and motifs are clustered based on their enrichment amongst proteins. The difference to the second heatmap is that the filter that requires a motif to be in 5% of proteins (and more than one protein) is applied. Clustering subseqeunctly restricted to the 30 motifs with the greatest overall enrichment in the dataset. Proteins containing none of the motifs are not displayed.
