## Supplemental File 4 for "ProminTools: Shedding light on proteins of unknown function in biomineralization with user friendly tools illustrated using mollusc shell matrix protein sequences": Mann14_cl1_Rscript.R

```
#' ---
#' title: Overrepresented motifs in your proteins of interest
#' author: A tool by Alastair Skeffington hosted by Cyverse and making use of the motif-x motif finding engine
#' ---
#'<style>
#'  body {
#'    text-align: justify}
#'</style>


#+ include = FALSE
library (wordcloud)
library(ggplot2)
library(ggrepel)
library(gplots)
library(knitr)
library(kableExtra)
library(ComplexHeatmap)
library(Rfast)

#'<br>
#'The main outputs of the program are listed below:  
#+ echo = FALSE, warnings=FALSE, messages=FALSE

#'* Mann14_cl1 _bgmotifs.txt : counts of each of the overrepresented motifs in each of the proteins in the background sequence set  
#'* Mann14_cl1 _fgmotifs.txt : counts of each of the overrepresented motifs in each of the proteins in the foreground sequence set  
#'* Mann14_cl1 _fgenrich.txt : the enrichment of each of the overrepresented motifs in each of the proteins in the foreground sequence set with repspect to the background sequence set  
#'* Mann14_cl1 _motifsummary.txt : for each overrepresented motif, the enrichemnt in the forgraound sequnces relative to the background; the count of proteins in which it appears and the median count of the motif per  protein  
#'* Mann14_cl1 _motifs.html : A html summary of the analysis with some helpful plots  
#'* Mann14_cl1 _Rscript.R : The R script that generated the html output. This can be used as a basis to produce your own custom figures and plots in R  
#'* Mann14_cl1 _wordclouds.svg : An svg file of the wordclouds from the html report. This can be imported into Inkscape or other software to make publication ready figures  

#+ echo = FALSE, warnings=FALSE, messages=FALSE

#'<br>
#'The main outputs of the program are the data tables. For convenience these are processed to a html document with
#'some visualizations of the data. However one analysis pipeline can never be appropriate for every dataset,
#'so the R script used to generate these diagrams is also an output of the program. This is to allow you, or an
#'infromatically minded colleague to easily tweak the plots or to develop these analyses further in a way 
#'appropriate for your data.  
#'<br>
#'The table below shows the motifs that are significantly overrepresented in your protein set of interest (PSOI) compared
#'to the background proteome. In your dataset 3 overrepresented motifs were identified. If you removed the PSOI from the background proteome fasta file
#'which you used as an input, the you may have motifs that are infinitely enriched ('Inf') because they are present
#'in the PSOI but not in the background.
#+ echo = FALSE, fig.height=3, fig.width=6

mots.path = "/home/mpimp-golm.mpg.de/skeffington/winhome/diatoms/cyverse_tools/motif_finder2/Mann14_cl1_motifsummary.txt"
mots_num.path  = "/home/mpimp-golm.mpg.de/skeffington/winhome/diatoms/cyverse_tools/motif_finder2/Mann14_cl1_fgmotifs.txt"
fg_enrich.path = "/home/mpimp-golm.mpg.de/skeffington/winhome/diatoms/cyverse_tools/motif_finder2/Mann14_cl1_fgenrich.txt"

wclname = "Mann14_cl1_wordcloud.svg"


mots<-read.table(mots.path, sep="\t", header=TRUE)
mot_num<-read.table(mots_num.path, sep="\t", header=TRUE, row.names=1)
fg_enrich<-read.table(fg_enrich.path, sep="\t", header=TRUE, row.names=1)

mot_num = mot_num[ rowSums(mot_num)!=0, ] 
protcount<-nrow(mot_num)

mots<-mots[order(mots$enrichment, decreasing = TRUE), ]
mots$enrichment<-round(mots$enrichment, 2)
if (nrow(mots)>12){
  kable(mots, row.names=FALSE, col.names=c("Motif", "Enrichment", "Count of proteins with motif", "Median  motif count per protein")) %>%
    kable_styling(bootstrap_options = c("striped", "hover", "condensed"), full_width = T) %>%
    column_spec(1, width = "5em") %>%
    column_spec(2, width = "6em") %>%
    column_spec(3, width = "15em") %>%
    column_spec(4, width = "18em") %>%
    scroll_box(width = "70%", height = "500px")
}else{
  kable(mots, row.names=FALSE, col.names=c("Motif", "Enrichment", "Count of proteins with motif", "Median  motif count per protein")) %>%
    kable_styling(bootstrap_options = c("striped", "hover", "condensed"), full_width = F, position = "left") %>%
    column_spec(1, width = "5em") %>%
    column_spec(2, width = "6em") %>%
    column_spec(3, width = "15em") %>%
    column_spec(4, width = "18em") 
}

#'<br>
#'The data in the table above can be visualized as wordclouds. To avoid plotting problems, any infinitely enriched
#'motifs were first removed from the data. In this analyses we are not interested in the peculiarities of 
#'particular proteins, but in the properties of this group of proteins as a whole, so any motifs found in less
#'than 5% of the proteins or in only one protein, were also removed before plotting.
#'
#'In the wordcoulds the height of the letters relates to either the number of proteins in the set of interest containing 
#'a given motif (left), the enrichment of a motif relative to the background proteome (middle), or the product of the scaled values of 
#'these two measures (right). In some datasets with very few motifs, the scaled values are numerically undefined and no wordcloud will be plotted.
#'

#+ echo = FALSE, warnings=FALSE, messages=FALSE
mots_finite<-subset(mots, enrichment!="Inf")
prots_5pc<-0.05*protcount
mots_finite<-subset(mots_finite, positiveprots >= prots_5pc & positiveprots > 1)

mots_finite$enrichment2<-mots_finite$enrichment
mots_finite$enrichment2[mots_finite$enrichment2 == 'Inf']<-1000000
mots_finite$enrichment2<-as.numeric(mots_finite$enrichment2)
mots_finite$enrichment_sca<- (mots_finite$enrichment2 - min(mots_finite$enrichment2)) / (max(mots_finite$enrichment2) - min(mots_finite$enrichment2))
mots_finite$positiveprots_sca<- (mots_finite$positiveprots - min(mots_finite$positiveprots)) / (max(mots_finite$positiveprots) - min(mots_finite$positiveprots))

mots_finite$scalprod<- (6 * mots_finite$enrichment_sca + 1) * (6 * mots_finite$positiveprots_sca + 1)

#+ echo = FALSE, warnings=FALSE, messages=FALSE, fig.height=3, fig.width=8
if(!anyNA(mots_finite$scalprod)){
	layout(matrix(c(1,2,3,4,5,6), nrow=2, byrow = TRUE), heights=c(0.7, 1.5))
	par(mar=rep(0, 4))
	plot.new()
	text(x=0.5, y=0.2, "Number of proteins\ncontaining motif", cex=1.5, pos=3)
	plot.new()
	text(x=0.5, y=0.2, "Enrichment", cex=1.5, pos=3)

	plot.new()
	text(x=0.5, y=0.2, "Product of scaled\nvalues", cex=1.5, pos=3)
	wordcloud(mots_finite$motif, mots_finite$positiveprots, scale=c(3, 0.2), random.order=FALSE, min.freq = 1, rot.per="0", colors= (brewer.pal(9, "YlOrRd")))

	wordcloud(mots_finite$motif, mots_finite$enrichment, scale=c(3, 0.2), min.freq = 1, random.order=FALSE, rot.per="0", colors= (brewer.pal(9, "YlOrRd")))

	wordcloud(mots_finite$motif, mots_finite$scalprod, scale=c(3, 0.2), random.order=FALSE, min.freq = 1, rot.per="0", colors= (brewer.pal(9, "YlOrRd")), main="product")
}

#+ echo = FALSE, message=FALSE, include=FALSE
if(!anyNA(mots_finite$scalprod)){
	svg(wclname)
	layout(matrix(c(1,2,3,4,5,6), nrow=2, byrow = TRUE), heights=c(0.7, 1.5))
	par(mar=rep(0, 4))
	plot.new()
	text(x=0.5, y=0.2, "Number of proteins\ncontaining motif", cex=1.5, pos=3)
	plot.new()
	text(x=0.5, y=0.2, "Enrichment", cex=1.5, pos=3)

	plot.new()
	text(x=0.5, y=0.2, "Product of scaled\nvalues", cex=1.5, pos=3)
	wordcloud(mots_finite$motif, mots_finite$positiveprots, scale=c(3, 0.2), random.order=FALSE, min.freq = 1, rot.per="0", colors= (brewer.pal(9, "YlOrRd")))

	wordcloud(mots_finite$motif, mots_finite$enrichment, scale=c(3, 0.2), min.freq = 1, random.order=FALSE, rot.per="0", colors= (brewer.pal(9, "YlOrRd")))

	wordcloud(mots_finite$motif, mots_finite$scalprod, scale=c(3, 0.2), random.order=FALSE, min.freq = 1, rot.per="0", colors= (brewer.pal(9, "YlOrRd")), main="product")

	dev.off()
}

#'<br>
#'In general there tends to be a negative correlation between the enrichment of a motif and the number of
#'proteins in which it is found. It is possible that motifs that deviate from this trend (ie they are 
#'unusually enriched given the number of proteins in which they are found, or are in an unusually 
#'large number of proteins given their enrichment) might have particular biological significance. The scatter
#'plots below plot the enrichment of the motifs against the number of proteins in which they are found (and vice versa)
#'to help you visualize any deviations from the expected negative correlation. If a linear regression is appropriate, a regression line (either linear or polynomial) is 
#'shown in red and the shaded area represents the 95% confidence interval of the regression. No regression line is shown in cases with strong non-linearity.
#'Points are labelled only if there is sufficient space in the plot.
#'<br>
#+
#+ echo = FALSE, warnings=FALSE, messages=FALSE
enrich_pos.lm<-lm(mots_finite$enrichment ~ mots_finite$positiveprots)
lmr2<-summary(enrich_pos.lm)$r.squared
if(lmr2 > 0.95){
  method_fit<-TRUE
  }else{
  method_fit<-FALSE}

#+ echo = FALSE, warnings=FALSE, messages=FALSE

if(nrow(mots_finite) < 40 ){
  scat1<-ggplot(mots_finite, aes(enrichment, positiveprots))
  scat1<-scat1 + geom_point() + theme_classic() + 
    {if(method_fit)geom_smooth(method = "lm", se = TRUE, show.legend = FALSE, aes(colour = "red"), weight = "1", fill = "#E5D0D0")} +
    geom_text_repel(aes(label = mots_finite$motif), size = 3.5) +
    labs(x="Fold motif enrichment", y = "Number of proteins containing motif")
  suppressWarnings(suppressMessages(print(scat1)))
}else{
  
    scat1<-ggplot(mots_finite, aes(enrichment, positiveprots))
    scat1<- scat1 + geom_point() + theme_classic() + 
    {if(method_fit)geom_smooth(method = "lm", se = TRUE, show.legend = FALSE, aes(colour = "red"), weight = "1", fill = "#E5D0D0")} +
    labs(x="Fold motif enrichment", y = "Number of proteins containing motif")
    suppressWarnings(suppressMessages(print(scat1)))
}


#'<br>
#+ echo = FALSE, warnings=FALSE, messages=FALSE

if(nrow(mots_finite) < 40 ){
  scat2<-ggplot(mots_finite, aes( positiveprots, enrichment))
  scat2<-scat2 + geom_point() + theme_classic() + 
    {if(method_fit)geom_smooth(method = "lm", se = TRUE, show.legend = FALSE, aes(colour = "red"), weight = "1", fill = "#E5D0D0")} +
    geom_text_repel(aes(label = mots_finite$motif), size = 3.5) +
    labs(y="Fold motif enrichment", x = "Number of proteins containing motif")
  suppressWarnings(suppressMessages(print(scat2)))
}else{
  scat2<-ggplot(mots_finite, aes( positiveprots, enrichment))
  scat2<-scat2 + geom_point() + theme_classic() + 
    {if(method_fit)geom_smooth(method = "lm", se = TRUE, show.legend = FALSE, aes(colour = "red"), weight = "1", fill = "#E5D0D0")} +
    labs(y="Fold motif enrichment", x = "Number of proteins containing motif")
  suppressWarnings(suppressMessages(print(scat2)))
        
}


#'<br>
#'You may be interested in grouping or subdividing your PSOI based on the motif content of the proteins. 
#'The heatmaps below provide a starting point for such approaches. The top heatmap displays a hierarchical
#'clustering of the proteins based on the number of motifs they contain, and simultaneously a 
#'clustering of motifs based on their distributions amongst proteins. In contrast to the above analyses,
#'the filters that remove the infinitely enriched motifs and motifs found in less than 5% of proteins 
#'are not applied. For clarity however, only the 30 motifs found most frequently amongst the proteins 
#'are used in the clustering for the first heatmap. Proteins containing none of the motifs are not displayed.
  
#+ echo=FALSE

hmcol <- colorRampPalette(brewer.pal(9, "GnBu"), bias = 2)(100)

mot_num <- as.matrix(mot_num)
mot_num<-mot_num[,order(colSums(mot_num), decreasing = TRUE)]
if(ncol(mot_num) > 30){
  mot_num<-mot_num[ , 1:30]
}

mot_num = mot_num[ rowSums(mot_num)!=0, ] 

heat1height<-0.22*nrow(mot_num) + 13
heat1width<-0.27*ncol(mot_num) + 10.2

addnoise2vec<- function(m){
  minvec<- m %>% rowMins()
  maxvec<- m %>% rowMaxs()
  Zerocorr<-0
  for(i in 1 : length(maxvec)){
    if(minvec[i] == maxvec[i]){
      Zerocorr <- 1
    }
  }
  if(Zerocorr == 1){
    matdim<-dim(m)[1] * dim(m)[2]
    m<- m + runif(m, min = 0, max = 0.0000005)
  }
  return(m)
}

mot_num<-addnoise2vec(mot_num)

#+ fig.height=heat1height, fig.width=heat1width, echo=FALSE, dpi = 50

HM_1<-Heatmap(mot_num, 
              col = hmcol,
              #clustering_distance_rows = "euclidean",
              clustering_distance_rows = function(x, y) 1 - cor(x, y),
              clustering_method_rows = "ward.D",
              clustering_distance_columns =  function(x, y) 1 - cor(x, y),
              clustering_method_columns = "ward.D",
              row_names_gp = gpar(fontsize = 14),
              column_names_gp = gpar(fontsize = 14),
              row_dend_width = unit(3, "cm"),
              column_dend_height = unit(3, "cm"),
              heatmap_legend_param = list(legend_width = unit(6, "cm"),
                                          title_position = 'topcenter',
                                          legend_direction = 'horizontal',
                                          title = "Motif count",
                                          title_gp = gpar(fontsize = 18),
                                          labels_gp = gpar(fontsize = 14)
              ))

draw(HM_1, heatmap_legend_side = "top")

#+

#+ echo = FALSE


mots_finite2<-subset(mots, enrichment!="Inf")
fg_enrich2 <- fg_enrich[, colnames(fg_enrich) %in% mots_finite2$motif]
fg_enrich2 <- as.matrix(fg_enrich2)

fg_enrich2<-fg_enrich2[,order(colSums(fg_enrich2),decreasing = TRUE)]
if(ncol(fg_enrich2) > 30){
  fg_enrich2<-fg_enrich2[ , 1:30]
}

fg_enrich2 = fg_enrich2[ rowSums(fg_enrich2)!=0, ] 

heat2height<-0.22*nrow(fg_enrich2) + 13
heat2width<-0.27*ncol(fg_enrich2) + 10.2
#+
#'<br>
#'In the second heatmap, proteins are clustered based on motif enrichment in a given protein sequence with 
#'respect the the background sequences, and motifs are clustered based on their enrichment amongst proteins.
#'The filter that removes motifs found in less than 5% of proteins is not applied, but infinitely enriched
#'motifs are excluded. However clustering is restricted to the 30 motifs with the greatest overall enrichment in 
#'the dataset. Proteins containing none of the motifs are not displayed.
#'<br>

#+ echo = FALSE

fg_enrich2<-addnoise2vec(fg_enrich2)

#+ fig.height=heat2height, fig.width=heat2width, echo=FALSE, dpi = 50


HM_2<-Heatmap(fg_enrich2, 
        col = hmcol,
        #clustering_distance_rows = "euclidean",
        clustering_distance_rows = function(x, y) 1 - cor(x, y),
        clustering_method_rows = "ward.D",
        clustering_distance_columns =  function(x, y) 1 - cor(x, y),
        clustering_method_columns = "ward.D",
        row_names_gp = gpar(fontsize = 14),
        column_names_gp = gpar(fontsize = 14),
        row_dend_width = unit(3, "cm"),
        column_dend_height = unit(3, "cm"),
        heatmap_legend_param = list(legend_width = unit(12, "cm"),
                                    title_position = 'topcenter',
                                    legend_direction = 'horizontal',
                                    title = "Fold motif enrichment",
                                    title_gp = gpar(fontsize = 18),
                                    labels_gp = gpar(fontsize = 14),
									labels_rot = 45
                                    ))

draw(HM_2, heatmap_legend_side = "top")


#+ echo = FALSE

fg_enrich3 <- fg_enrich[, colnames(fg_enrich) %in% mots_finite$motif]
fg_enrich3 <- as.matrix(fg_enrich3)
fg_enrich3<-fg_enrich3[,order(colSums(fg_enrich3),decreasing = TRUE)]
if(ncol(fg_enrich3) > 30){
  fg_enrich3<-fg_enrich3[ , 1:30]
}

fg_enrich3 = fg_enrich3[ rowSums(fg_enrich3)!=0, ] 

heat3height<-0.22*nrow(fg_enrich3) + 13
heat3width<-0.27*ncol(fg_enrich3) + 10.2
#+
#'<br>
#'In the third heatmap, proteins are clustered based on motif enrichment in a given protein sequence with 
#'respect the the background sequences, and motifs are clustered based on their enrichment amongst proteins.
#'The difference to the second heatmap is that the filter that requires a motif to be in 5% of proteins (and more than one protein) is 
#'applied. Clustering subseqeunctly restricted to the 30 motifs with the greatest overall enrichment in 
#'the dataset. Proteins containing none of the motifs are not displayed.
#'<br>

#+ echo = FALSE

fg_enrich3<-addnoise2vec(fg_enrich3)

#+ fig.height=heat3height, fig.width=heat3width, echo=FALSE, dpi = 50


HM_3<-Heatmap(fg_enrich3, 
              col = hmcol,
              #clustering_distance_rows = "euclidean",
              clustering_distance_rows = function(x, y) 1 - cor(x, y),
              clustering_method_rows = "ward.D",
              clustering_distance_columns =  function(x, y) 1 - cor(x, y),
              clustering_method_columns = "ward.D",
              row_names_gp = gpar(fontsize = 14),
              column_names_gp = gpar(fontsize = 14),
              row_dend_width = unit(3, "cm"),
              column_dend_height = unit(3, "cm"),
              heatmap_legend_param = list(legend_width = unit(12, "cm"),
                                          title_position = 'topcenter',
                                          legend_direction = 'horizontal',
                                          title = "Fold motif enrichment",
                                          title_gp = gpar(fontsize = 18),
                                          labels_gp = gpar(fontsize = 14),
										  labels_rot = 45
              ))

draw(HM_3, heatmap_legend_side = "top")
```
