## Supplemental File 5 for "ProminTools: Shedding light on proteins of unknown function in biomineralization with user friendly tools illustrated using mollusc shell matrix protein sequences"

#####################
The motif-x software:
#####################

Wagih O, Sugiyama N, Ishihama Y, Beltrao P. (2015) Uncovering phosphorylation-based specificities through functional interaction networks (2015). Mol. Cell. Proteomics 
Chou MF & Schwartz D (2011). Biological sequence motif discovery using motif-x. Curr Protoc Bioinformatics. Chapter 13:Unit 13.15-24. doi:10.1002/0471250953.bi1315s35.

The motif-x software may be used freely by users from academic and non-profit organizations. Users from the commercial sector should contact Daniel Schwartz (daniel.schwartz(at)uconn.edu).

The software in this package is provided under the GPL-3 licence: https://www.gnu.org/licenses/gpl-3.0

############
fLPS
############
This is used to look for biases in amino acid content in both regions of the protein as well as at the whole protein level. fLPS is designed to discover compositionally biased regions in proteins: http://biology.mcgill.ca/faculty/harrison/flps.html

Harrison, P.M. fLPS: Fast discovery of compositional biases for the protein universe. BMC Bioinformatics 18, 476 (2017) doi:10.1186/s12859-017-1906-3

Copyright 2017. Paul Martin Harrison.

Distrubuted under a 3-clause BSD license:

Redistribution and use in source and binary forms, with or without modification, are permitted provided that the following conditions are met:

Redistributions of source code must retain the above copyright notice, this list of conditions and the following disclaimer.

Redistributions in binary form must reproduce the above copyright notice, this list of conditions and the following disclaimer in the documentation and/or other materials provided with the distribution.

Neither the name of the copyright holder nor the names of its contributors may be used to endorse or promote products derived from this software without specific prior written permission.

This software is provided by the copyright holders and contributors "as is" and any express or implied warranties, including, but not limited to, the implied warranties of merchantability and fitness for a particular purpose are disclaimed. In no event shall the
 copyright holder or contributors be liable for any direct, indirect, incidental, special, exemplary, or consequential damages (including, but not limited to, procurement of substitute goods or services; loss of use, data, or profits; or business interruption) however
 caused and on any theory of liability, whether in contract, strict liability, or tort (including negligence or otherwise) arising in any way out of the use of this software, even if advised of the possibility of such damage.

####
Seg
####
Seg divides seqeunces into low and high complexity regions. More information is available here: http://www.biology.wustl.edu/gcg/seg.html

Authors: John Wootton and Scott Federhen
National Library of Medicine National Institutes of Health Bethesda, Maryland, MD 20894, USA

Seg was written by Wootton and Federhen at the National Center for Biotechnology Information (NCBI). Their public-domain program was modified by Scott Rose for distribution with Version 9 of the Wisconsin Package.

Wootton, J.C., Federhen, S. (1993) Statistics of local complexity in amino acid sequences and sequence databases. Computers & Chemistry 17: 149-163.

#####
VSL2
#####

VSL2 predicts regions of intrinsic disorder in proteins. Although not the most modern predictor available, it is still has good accuracy and is very fast compared to more recent predictors. This makes it feasible to run the predictor on a proteome scale in a reasonable timeframe.

Peng, K., Radivojac, P., Vucetic, S. et al. Length-dependent prediction of protein intrinsic disorder. BMC Bioinformatics 7, 208 (2006) doi:10.1186/1471-2105-7-208

Incorperated into ProminTools with the permission of Prof. Zoran Obradovic.

#####
SAPS
#####

SAPS calculates a variety of statistical properties for protein sequences.

Author: Volker Brendel Department of Mathematics, Stanford University, Stanford CA 94305, USA

Brendel, V., Bucher, P., Nourbakhsh, I.R., Blaisdell, B.E. & Karlin, S. (1992) Methods and algorithms for statistical analysis of protein sequences Proc. Natl. Acad. Sci. U.S.A. 89, 2002-2006.

Karlin, S., Weinstock, G.M., Brendel, V. (1995) Bacterial classifications derived from RecA protein sequence comparisons. J. Bacteriol. 177: 6881-6893.

Made freely available by the author. See http://brendelgroup.org/bioinformatics2go/bioinformatics2go.php

####
Perl
####

Perl is free software and is redistributed under the terms of the GNU General Public License. See https://perldoc.perl.org/perlgpl.html


###
R
###

R as a package is licensed under GPL-2 | GPL-3. 
See https://www.r-project.org/Licenses/
