## Supplemental File 2 for "ProminTools: Shedding light on proteins of unknown function in biomineralization with user friendly tools illustrated using mollusc shell matrix protein sequences": Mann2014all_motifs.html

Overrepresented motifs in your proteins of interest


### Overrepresented motifs in your proteins of interest

###### A tool by Alastair Skeffington hosted by Cyverse and making use of the motif-x motif finding engine

#### 2020-02-26

The main outputs of the program are listed below:

- Mann2014all \_bgmotifs.txt : counts of each of the overrepresented motifs in each of the proteins in the background sequence set
- Mann2014all \_fgmotifs.txt : counts of each of the overrepresented motifs in each of the proteins in the foreground sequence set
- Mann2014all \_fgenrich.txt : the enrichment of each of the overrepresented motifs in each of the proteins in the foreground sequence set with repspect to the background sequence set
- Mann2014all \_motifsummary.txt : for each overrepresented motif, the enrichemnt in the forgraound sequnces relative to the background; the count of proteins in which it appears and the median count of the motif per protein
- Mann2014all \_motifs.html : A html summary of the analysis with some helpful plots
- Mann2014all \_Rscript.R : The R script that generated the html output. This can be used as a basis to produce your own custom figures and plots in R
- Mann2014all \_wordclouds.svg : An svg file of the wordclouds from the html report. This can be imported into Inkscape or other software to make publication ready figures

The main outputs of the program are the data tables. For convenience these are processed to a html document with some visualizations of the data. However one analysis pipeline can never be appropriate for every dataset, so the R script used to generate these diagrams is also an output of the program. This is to allow you, or an infromatically minded colleague to easily tweak the plots or to develop these analyses further in a way appropriate for your data.

| Motif | Enrichment | Count of proteins with motif | Median motif count per protein |
| --- | --- | --- | --- |
| D.ID.IND | Inf | 1 | 0 |
| D.MN.GQ | Inf | 1 | 0 |
| DE.AD.ID | Inf | 1 | 0 |
| DM.NMN.G | Inf | 1 | 0 |
| DNID.IN | Inf | 1 | 0 |
| G.FGPGG | Inf | 1 | 0 |
| G.G.DLGG | Inf | 1 | 0 |
| GG.MD.GG | Inf | 1 | 0 |
| GPGGPG..G | Inf | 1 | 0 |
| GPGGPGGP | Inf | 1 | 0 |
| GQ.GP..M | Inf | 1 | 0 |
| IDEI.DE | Inf | 1 | 0 |
| M.NGQ.G | Inf | 1 | 0 |
| NAD.ID.I | Inf | 1 | 0 |
| NGQ.G..DM | Inf | 1 | 0 |
| PEV.P.V.P | Inf | 1 | 0 |
| PGGPGG.G | Inf | 1 | 0 |
| Q…TAQ.G | Inf | 1 | 0 |
| TPGVEP | Inf | 1 | 0 |
| V.PEV.P.V | Inf | 1 | 0 |
| VTP.V.PE | Inf | 1 | 0 |
| GPGG.GG | 182.69 | 2 | 0 |
| QA.PG..QA | 87.69 | 1 | 0 |
| TNQA.P | 81.11 | 1 | 0 |
| GGPGG | 80.77 | 5 | 0 |
| TN..H.GT | 79.17 | 1 | 0 |
| P.TNQA | 78.92 | 1 | 0 |
| NQA.P.T.Q | 78.30 | 1 | 0 |
| QAYP..N | 78.30 | 1 | 0 |
| PGTN.A | 76.73 | 1 | 0 |
| GQLG.R | 71.67 | 1 | 0 |
| FTGQ.G | 61.66 | 1 | 0 |
| QP..QP.A | 61.66 | 5 | 0 |
| F..QSNQ | 61.07 | 1 | 0 |
| P.VE.N.Q | 60.03 | 2 | 0 |
| QPPVE | 59.79 | 3 | 0 |
| NQQF..Q | 57.77 | 1 | 0 |
| P..NQA.P | 57.69 | 2 | 0 |
| QQP.VE | 57.42 | 2 | 0 |
| QQP..EL | 57.30 | 3 | 0 |
| F…LG.RG | 54.81 | 1 | 0 |
| N.Q.N.Q.N | 54.81 | 1 | 0 |
| P.TG.QP | 54.81 | 1 | 0 |
| Q.NQQF | 54.81 | 1 | 0 |
| D.IDT..D | 51.58 | 1 | 0 |
| Q.N.Q.N.Q | 50.75 | 1 | 0 |
| QQ.NS…Q | 49.33 | 2 | 0 |
| RG.D..GF | 48.72 | 1 | 0 |
| E.N.QQP | 48.23 | 2 | 0 |
| Q.PP.QP | 44.37 | 4 | 0 |
| C.CK..P | 43.06 | 3 | 0 |
| TPPVV | 38.82 | 3 | 0 |
| T.PVV.Y | 37.37 | 1 | 0 |
| VNPG.E.P | 36.54 | 1 | 0 |
| G.GMG.G | 31.97 | 2 | 0 |
| P…KCC | 30.69 | 5 | 0 |
| NVNPG | 28.42 | 1 | 0 |
| Q.NQQ | 27.19 | 5 | 0 |
| C..D..CP | 25.96 | 5 | 0 |
| Q..YQP | 23.78 | 5 | 0 |
| P…N..QP | 23.59 | 7 | 0 |
| C…CG..C | 23.36 | 6 | 0 |
| Q.PAQ | 20.74 | 6 | 0 |
| G…D..GC | 19.82 | 6 | 0 |
| QQP…P | 18.75 | 13 | 0 |
| Q.QP..Q | 18.05 | 13 | 0 |
| G..GGAG | 17.78 | 5 | 0 |
| PAQP | 17.36 | 6 | 0 |
| QPP..Q | 15.35 | 7 | 0 |
| QQPP | 14.86 | 15 | 0 |
| QP..YP | 14.33 | 7 | 0 |
| Q..QQ…Q | 14.21 | 12 | 0 |
| QPR.Q | 12.18 | 7 | 0 |
| P..NP.P | 11.07 | 8 | 0 |
| NG.GG | 10.25 | 13 | 0 |
| P…P..QP | 10.10 | 11 | 0 |
| PV.P.P | 8.83 | 11 | 0 |
| QQP | 7.55 | 35 | 0 |
| QP..Q | 6.67 | 32 | 0 |
| Q.N.Q | 6.18 | 25 | 0 |
| G..G..GG | 6.00 | 23 | 0 |
| Q..QQ | 5.69 | 41 | 0 |
| Q.QP | 5.59 | 32 | 0 |
| G.G.GG | 5.48 | 25 | 0 |
| CCP | 5.33 | 18 | 0 |
| P.QQ | 5.28 | 28 | 0 |
| GGA.G | 5.21 | 11 | 0 |
| TTT..P | 4.91 | 19 | 0 |
| P.QP | 4.82 | 37 | 0 |
| P..QP | 4.53 | 36 | 0 |
| PQ…Q | 4.49 | 35 | 0 |
| Q..PQ | 4.34 | 35 | 0 |
| QP..P | 4.16 | 37 | 0 |
| PG..P | 4.14 | 49 | 0 |
| Q..QP | 4.03 | 31 | 0 |
| P..N.Q | 3.97 | 35 | 0 |
| D…D..DD | 3.91 | 13 | 0 |
| C…P.G | 3.87 | 23 | 0 |
| Q.G…Q | 3.86 | 33 | 0 |
| D..DD..D | 3.82 | 13 | 0 |
| QPT | 3.68 | 42 | 0 |
| Q.P.T | 3.52 | 45 | 0 |
| G..GG | 3.51 | 71 | 0 |
| NQ.P | 3.43 | 40 | 0 |
| AGQ | 3.42 | 48 | 0 |
| PA..P | 3.21 | 38 | 0 |
| P..NP | 2.95 | 42 | 0 |
| TTTT | 2.93 | 14 | 0 |
| CP.G | 2.82 | 30 | 0 |
| P.T.T | 2.82 | 62 | 0 |
| QP | 2.43 | 181 | 0 |
| TTP | 2.31 | 54 | 0 |
| Q.P | 2.26 | 167 | 0 |
| Q..P | 2.25 | 174 | 0 |
| P.Q | 2.06 | 182 | 0 |
| PG | 1.87 | 227 | 1 |
| CP | 1.87 | 120 | 0 |
| A.P | 1.83 | 207 | 1 |
| AP | 1.83 | 198 | 1 |
| P…C | 1.82 | 122 | 0 |
| GG | 1.80 | 270 | 1 |
| M..M | 1.78 | 91 | 0 |
| N.Q | 1.78 | 205 | 1 |
| G.P | 1.78 | 211 | 1 |
| P.G | 1.77 | 234 | 1 |
| P.P | 1.74 | 187 | 0 |
| GQ | 1.70 | 225 | 1 |
| Q.G | 1.70 | 218 | 1 |
| G.G | 1.68 | 266 | 1 |
| P..G | 1.68 | 202 | 1 |
| Q…G | 1.67 | 208 | 1 |
| A…P | 1.66 | 212 | 1 |
| P..N | 1.66 | 194 | 1 |
| MP | 1.64 | 130 | 0 |
| AQ | 1.64 | 214 | 1 |
| G…D | 1.60 | 264 | 1 |
| T…P | 1.60 | 207 | 1 |
| N.G | 1.55 | 230 | 1 |
| M…G | 1.55 | 168 | 0 |
| G.N | 1.55 | 247 | 1 |
| P.V | 1.54 | 224 | 1 |
| G.Q | 1.54 | 207 | 1 |
| V.P | 1.53 | 209 | 1 |
| NG | 1.53 | 249 | 1 |
| V…P | 1.51 | 231 | 1 |
| T…A | 1.49 | 246 | 1 |
| NP | 1.49 | 198 | 1 |
| A…A | 1.49 | 247 | 1 |
| P.N | 1.48 | 194 | 1 |
| P..T | 1.48 | 206 | 1 |
| T.P | 1.48 | 224 | 1 |
| A…G | 1.47 | 273 | 1 |
| Y.P | 1.45 | 178 | 0 |
| P..V | 1.44 | 228 | 1 |
| T…G | 1.44 | 241 | 1 |
| G..T | 1.43 | 257 | 1 |
| RG | 1.43 | 235 | 1 |
| G..D | 1.43 | 234 | 1 |
| A..T | 1.42 | 255 | 1 |
| G…V | 1.42 | 273 | 1 |
| G…Y | 1.42 | 196 | 1 |
| A..D | 1.41 | 229 | 1 |
| NA | 1.40 | 227 | 1 |
| W.G | 1.40 | 79 | 0 |
| S…C | 1.39 | 143 | 0 |
| V.G | 1.39 | 286 | 1 |
| Y…G | 1.38 | 206 | 1 |
| R.P | 1.37 | 171 | 0 |
| PR | 1.37 | 178 | 0 |
| DC | 1.37 | 117 | 0 |
| DP | 1.37 | 201 | 1 |
| GS | 1.36 | 264 | 1 |
| G.Y | 1.36 | 195 | 1 |
| S…G | 1.36 | 260 | 1 |
| G..R | 1.36 | 213 | 1 |
| SG | 1.35 | 289 | 2 |
| DA | 1.35 | 238 | 1 |
| N.P | 1.34 | 200 | 1 |
| C..K | 1.34 | 121 | 0 |
| S.Q | 1.34 | 218 | 1 |
| A.N | 1.34 | 233 | 1 |
| R..P | 1.33 | 189 | 0 |
| S..G | 1.33 | 284 | 1 |
| P…R | 1.33 | 188 | 0 |
| S…Q | 1.33 | 224 | 1 |
| F…G | 1.33 | 222 | 1 |
| D…D | 1.32 | 225 | 1 |
| PS | 1.31 | 241 | 1 |
| G…S | 1.31 | 282 | 1 |
| P..E | 1.30 | 215 | 1 |
| G..V | 1.30 | 267 | 1 |
| YD | 1.30 | 190 | 0 |
| DG | 1.29 | 254 | 1 |
| D.R | 1.28 | 213 | 1 |
| K..C | 1.28 | 129 | 0 |
| T..T | 1.28 | 223 | 1 |
| S.G | 1.28 | 268 | 1 |
| P…S | 1.27 | 218 | 1 |
| G…E | 1.27 | 241 | 1 |
| G.E | 1.27 | 224 | 1 |
| KC | 1.26 | 148 | 0 |
| K…C | 1.25 | 137 | 0 |
| P…I | 1.23 | 205 | 1 |
| A..E | 1.23 | 240 | 1 |
| G..F | 1.22 | 218 | 1 |
| E..G | 1.21 | 231 | 1 |
| K..A | 1.21 | 239 | 1 |
| V.V | 1.20 | 263 | 1 |
| K…G | 1.18 | 239 | 1 |
| PI | 1.18 | 201 | 1 |
| GF | 1.18 | 218 | 1 |
| S.T | 1.17 | 252 | 1 |
| GK | 1.17 | 267 | 1 |
| G..K | 1.16 | 238 | 1 |
| IG | 1.15 | 248 | 1 |
| GR | 1.15 | 226 | 1 |
| I.P | 1.15 | 211 | 1 |
| G.K | 1.13 | 256 | 1 |
| L..G | 1.13 | 283 | 1 |
| GL | 1.13 | 281 | 1 |
| G…I | 1.12 | 246 | 1 |
| LG | 1.12 | 275 | 1 |
| A..K | 1.11 | 256 | 1 |
| L..P | 1.11 | 232 | 1 |
| I.A | 1.10 | 243 | 1 |
| G..L | 1.09 | 276 | 1 |
| I.D | 1.08 | 245 | 1 |
| D…I | 1.07 | 232 | 1 |
| D..K | 1.02 | 253 | 1 |
